# Differential CD8+ T/NK cell-mediated reduction of HIV-1 replication after combination of ART with TIGIT or KLRG1 blockade in humanized mice

**DOI:** 10.1101/2025.09.11.675307

**Authors:** I Sánchez-Cerrillo, I Tsukalov, M Agudo-Lera, O Popova, P Fuentes, J Alcain, R Gonzalez, L García-Fraile, I de los Santos, M Lázaro-Diez, D Perea, J Grau-Expósito, J Sevilla-Montero, Vladimir Vrbanac, Alejandro Balazs, C Muñoz-Calleja, M.L Toribio, F Sánchez-Madrid, JG Prado, M Genescà, MJ Buzón, E Martín-Gayo

## Abstract

Expression of TIGIT and KLRG1 has been associated to an exhausted, dysfunctional state in natural killer (NK) and CD8+ T cells from people with HIV-1 (PWH), limiting the efficacy of immunotherapies aiming at achieving a functional cure of the infection. Antiretroviral therapy (ART) does not completely reverse this immune exhaustion, and its combination with blockade of immune checkpoint receptors such as TIGIT and KLRG1 could be a promising strategy to promote control of viral replication in PWH. However, the impact of targeting these two immune checkpoint receptors has not been evaluated *in vivo*. In this study, we used a humanized Bone Marrow, Liver and Thymus (hBLT) mouse model of HIV-1 infection to study the impact of ART in combination with aTIGIT or aKLRG1 or a bispecific aTIGIT/aKLRG1 mAbs. Our results indicated that combination of ART with either aTIGIT or aKLRG1 mAbs led to faster reduction of HIV-1 pVL. Furthermore, viral rebound after ART interruption (ATI) was delayed in mice treated with aTIGIT and aKLRG1 mAbs. Histological detection of HIV-1 p24 in the spleen was restricted to the white pulp in the aKLRG1 mAb-treated group, which correlated with higher infiltration of IFNγ+ CD8+ T cells in these histological regions and with increased cytotoxic CD107a+ Granzyme B+ CD8+ T cells in the spleen. In contrast, control of HIV-1 replication induced by the aTIGIT mAb was associated with an increased splenic CD107a+ IFNγ+ NKG2C+ CD57-adaptive NK cells. In contrast, combination of ART with a bispecific aTIGIT/aKLRG1 mAb was unable to efficiently suppress viral replication or delay viral rebound after ATI, potentially by inducing apoptosis of adaptive NKG2C+ NK. Together, these results suggest that combination of ART with individual TIGIT or KLRG1 blockade may be a promising immunotherapy strategy against HIV-1 by eliciting differential immune control mechanisms.

## INTRODUCTION

Antiretroviral therapy (ART) efficiently suppresses viral replication leading to undetectable levels of plasma viremia in people with HIV-1 (PWH), thus preventing transmission to other individuals. However, ART does not cure the infection, as the virus persists in a latent proviral form integrated in the DNA of target cells, mainly memory CD4+ T cells ^1–3^, which are predominantly located in different tissues, including lymphoid organs and the gastrointestinal associated lymphoid tissues (GALT) ^4–8^. A fraction of these latently infected T cells contains intact viral sequences and can produce replication-competent HIV-1 particles in the absence of ART ^9,10^. One of the HIV-1 cure strategies designed to eliminate persistent reservoirs that continues to be tested is the “shock and kill” approach, which involves the reactivation of HIV-1 replication using latency-reversing agents (LRAs), followed by clearance of the infected cells by stimulating the antiviral immune response ^5,11^. However, tested strategies based on LRAs alone have not achieved a complete reduction of the viral reservoir, despite successful viral reactivation *in vitro* ^12^. Therefore, it is essential to potentiate an immune response that efficiently depletes cells capable of producing new viral particles ^13^.

Chronic HIV-1 infection induces continuous immune system stimulation potentially due to sporadic viral reactivations ^14^, which contributes to the development of exhaustion in immune cell subtypes key for viral control, such as CD8+ T lymphocytes and Natural Killer (NK) cells. Exhausted immune cells are characterized by the expression of multiple inhibitory immune checkpoint receptors, leading to a dysfunctional state and the inability to eliminate infected cells ^15–17^. Currently, the dysfunctional and exhausted state of immune cells and the expression of checkpoint receptors do not appear to be fully reversed following long-term ART treatment ^18–21^.

Therefore, new strategies are needed to restore the function of exhausted CD8+ T cells and NK cells, in order to enhance elimination of persistent HIV-1-infected cells, and ultimately achieve a functional cure. An attractive possibility is the blockade of inhibitory receptors using specific monoclonal antibodies (mAb) as an immunotherapy ^22^. Among different potential candidate targets to reverse immune exhaustion, PD-1 represents the inhibitory receptor most extensively studied in CD8+ T and NK cells in different chronic diseases including cancer and HIV-1 infection ^23,24^. PD-1 modulation has shown promise in enhancing viral clearance in the context of HIV-1 infection ^18,25–27^. More recently, the checkpoint molecule T cell immunoreceptor with Ig and ITIM domains (TIGIT) has emerged as a key target for improving the function of both NK cells and CD8+ T cells against HIV-1-infected cells alone or in combination with activated dendritic cells (DCs) ^18,28–30^. In addition, the Killer cell Lectin-like Receptor G1 (KLRG1) molecule also expressed by both NK cells and CD8+ T cells has recently been proposed as a potential therapeutic target to enhance NK cell function and their ability to eliminate HIV-1-infected cells ^31,32^. Specifically, in CD8+ T cells, KLRG1 expression has been associated with control of other chronic viral infections; however, in the context of HIV-1, its role in this T cell population has not been well studied ^33,34^.

Most of these immunotherapeutic approaches have been tested only *in vitro*, but they have not yet been compared on *in vivo* models of HIV-1 infection. Moreover, most immunotherapies are administered systemically, and it is unclear which immune cell populations may be preferential affected and the impact on the antiviral immune response by targeting different immune checkpoint receptors in cells located in tissues. In addition, a potential beneficial strategy to boost the immune responses is the use of bispecific antibodies directed to different receptors ^35^, but it is unclear whether this strategy could be beneficial promoting protective antiviral immunity in lymphoid tissue. Therefore, to evaluate these questions it is critical the use of *in vivo* animal models of HIV-1 infection.

The non-human primate models (NHP) of HIV-1 infection with the simian immunodeficiency virus (SIV), have extensively employed for the development of vaccines and immunotherapies against HIV-1 ^22,24,36–38^. However, previous studies have underscored the limitations of NHP models due to genetic differences between the host (rhesus macaques versus humans) and the pathogen, which often lack epitopes similar to those found in the human virus, highlighting the need of a model that allows to study the response of human immune cells. The humanized bone marrow–liver–thymus (hBLT) mouse was developed several years ago as an alternative modeling of HIV-1 infection and human immunity *in vivo* ^39–43^. This model recapitulates seemingly the entire human immune system, including the presence of an autologous human thymus which enables the development of robust polyclonal human T cell responses against HIV-1 after infection ^44^ as well as some aspects of the disease progression, such as increase of plasma viral loads, the depletion of CD4+ T cells and the stimulation of HIV-1 specific T cells and viral evasion ^45^. Furthermore, viral suppression can be achieved in HIV-1 infected hBLT mice by administering oral ART, leading to an induced viral suppression state that closely resembles chronic infection present in PWH on ART ^46–48^. In addition, our group and others have described the expression of checkpoint receptors such as TIGIT and KLRG1 in NK cells from PWH and HIV-1 infected hBLT mice even after ART initiation ^21,28,31^. Thus, this model may serve as a valuable tool for studying chronic HIV-1 infection, including the association of exhaustion markers with viral replication and evaluating the impact of immunotherapeutic strategies targeting these receptors to modulate antiviral immune responses. One of the limitations of hBLT mice is the poor reconstitution of NK cells due to lack of homology between human and murine IL-15 ^49^. However, exogenous administration of human IL-15 can be used to overcome this issue and enhance NK cell engraftment and maturation ^50^, optimizing this animal model to test NK cell-based immunotherapies.

In this study, we evaluated the impact of individual or bispecific antibodies targeting the checkpoint receptors TIGIT and KLRG1 in combination with ART on promoting immune control of HIV-1 replication through activation of NK and/or CD8+ T cells responses in the hBLT mouse model. Mice treated with either individual aTIGIT or aKLRG1 mAb at ART initiation showed an earlier reduction of HIV-1 viral loads. Moreover, HIV-1 viral rebound was significantly delayed after ART interruption (ATI) in the presence of aTIGIT or aKLRG1 mAbs. Notably, differential immune mechanisms associated with either a specific enrichment in cytotoxic CD107a+ adaptive NKG2C+ NK cells responses or cytotoxic CD107a+ Granzyme B+ CD8 T cells was associated with enhanced HIV-1 control after the aTIGIT or aKLRG1 mAb treatment, respectively. Strikingly, HIV-1 suppression at ART initiation and control after ATI was impaired in hBLT mice injected with bispecific aTIGIT/aKLRG1 mAb, which appeared to specifically induce apoptosis of NK cells and reduce levels of CD8+ T cell activation *in vitro*. Together, these data suggest that blockade of TIGIT or KLRG1 in combination with ART could control of HIV-1 replication in the hBLT model by eliciting differential immune response mechanisms that may be useful for future personalized functional cure strategies.

## RESULTS

### Individual and bispecific targeting of TIGIT and KLRG1 differentially increase activation of CD56+ CD16+ NK cells and CD8+ T cells in vitro

MAbs recognizing TIGIT (Tiragolumab) and KLRG1 (HG1N07) were selected for our study based on previous *in vitro* or *in vivo* evidence of efficacy (Ghasemi, 2025). Additionally, a bispecific Ab containing the Fab regions from each of those individual mAbs and of the same isotype was also developed for this study (Figure 1A). First, the binding of these antibodies to different immune cells, such as NK cells or CD8+ T cells, was tested in peripheral blood mononuclear cells (PBMCs) from PWH *in vitro* (Supplemental Figure 1 A-B). Using a fluorochrome-labeled anti-human IgG (hIgG) secondary Ab, an increase in proportions of hIgG-labelled CD3+ T and CD56+CD16+ NK cells from PBMCs was observed when cells were cultured for 16h in the presence of aTIGIT, aKLRG1 or the bispecific mAbs, compared with control cells cultured only with secondary hIgG Ab (Supplemental Figure 1B, C).

**Figure 1.**
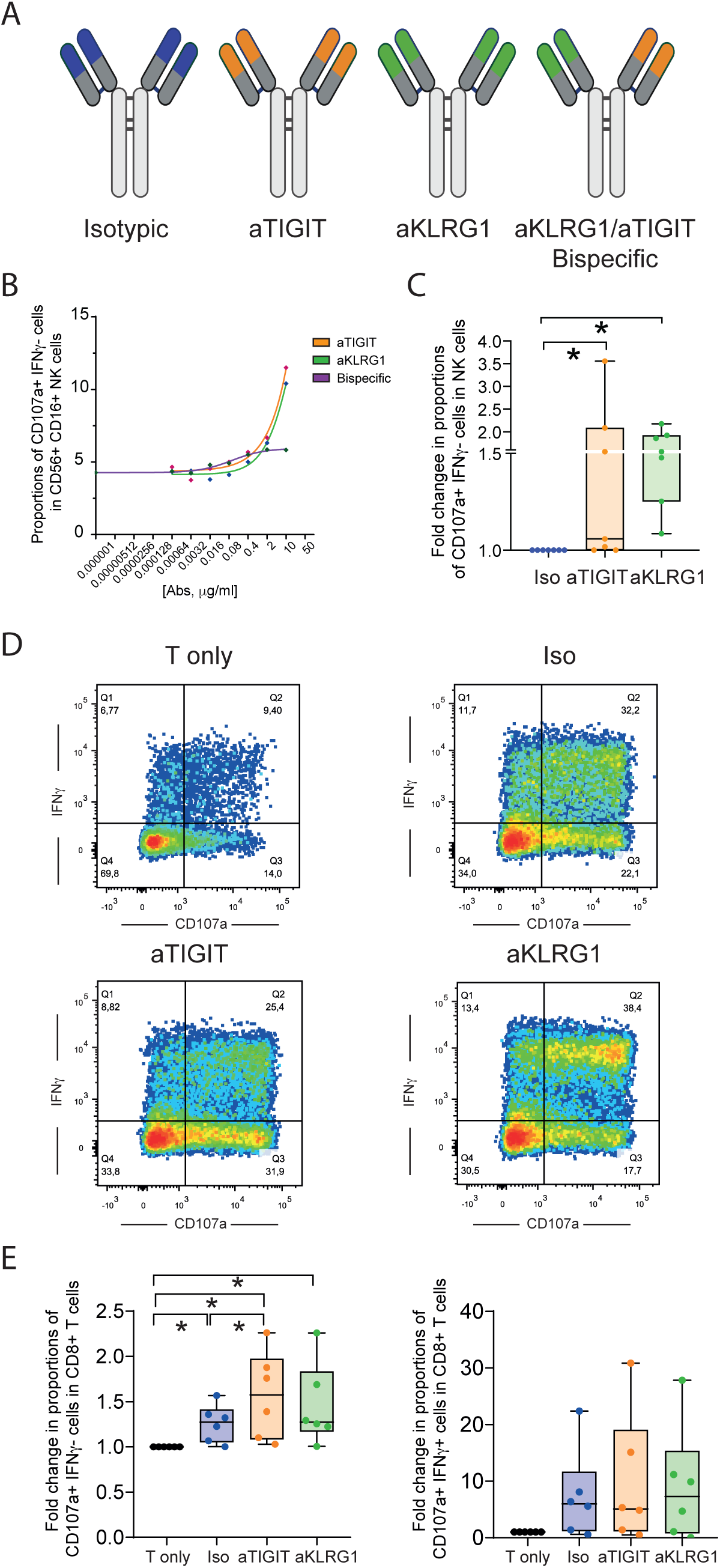
*In vitro* functional analysis of individual and bispecific antibodies directed to TIGIT and/or KLRG1 in NK and CD8+ T cells. (A): Representative schematic representation of the different antibodies used in the study. Variable chain regions are highlighted in green for those recognizing KLRG1 and in orange for those recognizing TIGIT. The human IgG1 heavy chain region was consistent across all antibodies. (B): Dose response analysis of intracellular expression of CD107a and IFNγ in NK from a selected PWH after culture with increasing concentrations of aTIGIT (orange), aKLRG1 (green) and bispecific (purple) mAbs. (C): Fold change in proportions to isotype treated-group of CD107a+ IFNγ-cells within CD56+ CD16+ NK cells from n=7 PWH, in response to the selected 5μg/ml of the different antibody treatment groups. (D): Representative flow cytometry plots showing IFNγ versus CD107a expression in CD8+ T cells after MLR assay under different individual or bispecific aTIGIT and aKLRG1 mAbs. (E): Fold change in proportions of CD107a+ IFNγ-(left) and CD107a+ IFNγ+ (right) in CD8+ T cells from n=6 HIV negative donors, in the presence of the different mentioned mAbs. In panel C and E, data are presented as Box and Whiskers plots showing median values, with minimum and maximum error bars. Statistical significance was calculated using a using a Wilcoxon t test. *p<0.05.

Next, we assessed function of NK and CD8+ T cells after incubation with the different mAbs in different functional assays. Different doses of aTIGIT, aKLR1 and the bispecific mAbs were tested for the induction of degranulation marker (CD107a) and IFNγ secretion by human NK cells from a PWH donor in the presence of susceptible K562 target cell line in a dose-response curve. Antibody concentrations between 2-10 μg/ml were found to be optimal, as higher proportions of CD107a+ IFNγ- and CD107a+ IFNγ+ cytotoxic cells were observed within CD56+ CD16+ NK cells treated with aTIGIT (orange) and aKLRG1 (green) mAbs (Figure 1B; Supplemental Figure 1D, E). In the case of bispecific mAb (shown in purple), saturation of induction of cytotoxic NK cells was observed at even lower concentrations of approximately 0.08 μg/mL (Figure 1B; Supplemental Figure 1E).

We then analyzed the expression of CD107a and IFNγ in NK cells from n=7 PWH cocultured for 4h in the presence of the K562 cell line using the optimal mAb concentration (Supplemental Figure 1D). Increased proportions of CD107a+ IFNγ- in CD56+ CD16+ NK cells were observed following TIGIT and KLRG1 blockade compared to cells incubated with the isotypic control mAb (Figure 1C). Additionally, a tendency for higher proportions of CD107a+ IFNγ+ in CD56+ CD16+ NK cells was detected after aTIGIT Ab treatment compared with aKLRG1 Ab (Supplemental Figure 1F). Therefore, individual or combined targeting of TIGIT and KLRG1 induces activation of NK cells from PWH *in vitro*.

On the other hand, to evaluate the functional impact of these mAbs in the activation and induction of cytotoxic CD8+ T cells, a mixed lymphocyte reaction (MLR) assay was used by culturing T lymphocytes isolated from healthy donors with allogenic primary dendritic cells (DCs), which allows to analyze polyclonal T cell proliferation and expression of cytotoxic and activation markers such as IFNγ, TNFα and CD107a (Figure 1D, Supplemental Figure 2A, B). In these experiments, significantly higher proportions of CD107a+ IFNγ- were observed in CD8+ T cells cultured with allogeneic DC for 6-7 days in the presence of the aTIGIT mAb compared to the isotype control or T cells cultured in media alone (Figure 1D-E). A trend toward increased proportions of CD107a+ IFNγ+ in CD8+ T was detected in response to allogeneic DC and aKLRG1 antibody treatment (Figure 1E, right). Regarding CD107a+ TNFα+ CD8+ T cells, a tendency for increase was also detected after aKLRG1 mAb incubation (Supplemental Figure 2C). In contrast, the presence of the bispecific mAb led to lower proportions of CD107a+ IFNγ-, CD107a+ IFNγ+ and CD107a+ TNFα+ CD8+ T cells compared with cells cultured in the presence of the isotypic control mAb (Supplemental Figure 2D). Together, the data suggest that individual and bispecific aTIGIT/aKLRG1 mAbs differentially affect the activation of NK and CD8+ T cells.

### Combination of ART and aTIGIT or aKLRG1 mAb immunotherapy promotes faster viral suppression and control HIV-1 replication after ATI

Next, we assessed the capacity of individual and bispecific immunotherapeutic targeting of TIGIT and KLRG1 to induce viral suppression after HIV-1 infection in the presence of ART and potentially control of viral replication after ATI using the hBLT mouse model ^40,52,53^. While hBLT mice can develop human NK cells, the maturation and functionality of these cells is less effective due to the lack of homology between human and murine IL-15 ^54^. In our previous studies, we optimized the reconstitution of human NK cells prior to infection by injection of rhIL-15 ^28^. This rh-IL15 administration protocol was also applied for two weeks prior to HIV-1 infection, when a significant increase of human CD56+ NK cell reconstitution was reached, which was also accompanied by a significant increase in human hCD45+ cells, CD4+ and CD8+ T cell numbers (Supplemental Figure 3A-B-C). Reconstituted hBLT mice were then infected intravenously (IV) with a high dose of the HIV-1 JRCSF strain (5,000 TCID_50_) and remained untreated for two weeks to allow HIV-1 replication (Supplemental Figure 3D). Since ART can be administered to these mice orally, we initiated oral ART at 2 weeks post-infection in different groups of mice simultaneously receiving either an isotypic IgG1 or individual aTIGIT or aKLRG1 or bispecific aTIGIT/aKLRG1 mAbs at treatment initiation and during ART maintenance for 11 days (Supplemental Figure 3D). Plasma viral loads (pVL) were assessed prior and after 7 and 11 days after ART initiation (8, 15 and 19 days post-infection, respectively) and we analyzed the kinetics of viral suppression in the different animal groups (Supplemental Figure 3D). Of note, no evident differences were observed between groups in terms of weekly weight measurements during the whole experiment (Supplemental Figure 3E). Prior to ART, the median levels of HIV-1 pVL at eight days post-infection oscillated from 10^4^-10^5^ copies/mL, and no statistical differences were detected between the different groups (Supplemental Figure 4A). Interestingly, while no significant changes were observed in the kinetics of pVL decay after ART initiation (Supplemental Figure 4B), a higher percentage of aviremic hBLT mice were observed already at day 7 post-ART initiation (day 15 post-infection) when aTIGIT (80%) and aKLRG1 (60%) mAb immunotherapies were combined at ART initiation, compared with animals receiving ART and isotype (40%) treatment (Figure 2A). Surprisingly, we observed a partial reduction of HIV-1 pVL in the presence of ART and only 20% of mice became aviremic in the group of hBLT mice treated with bispecific mAb (Supplemental Figure 4C).

**Figure 2.**
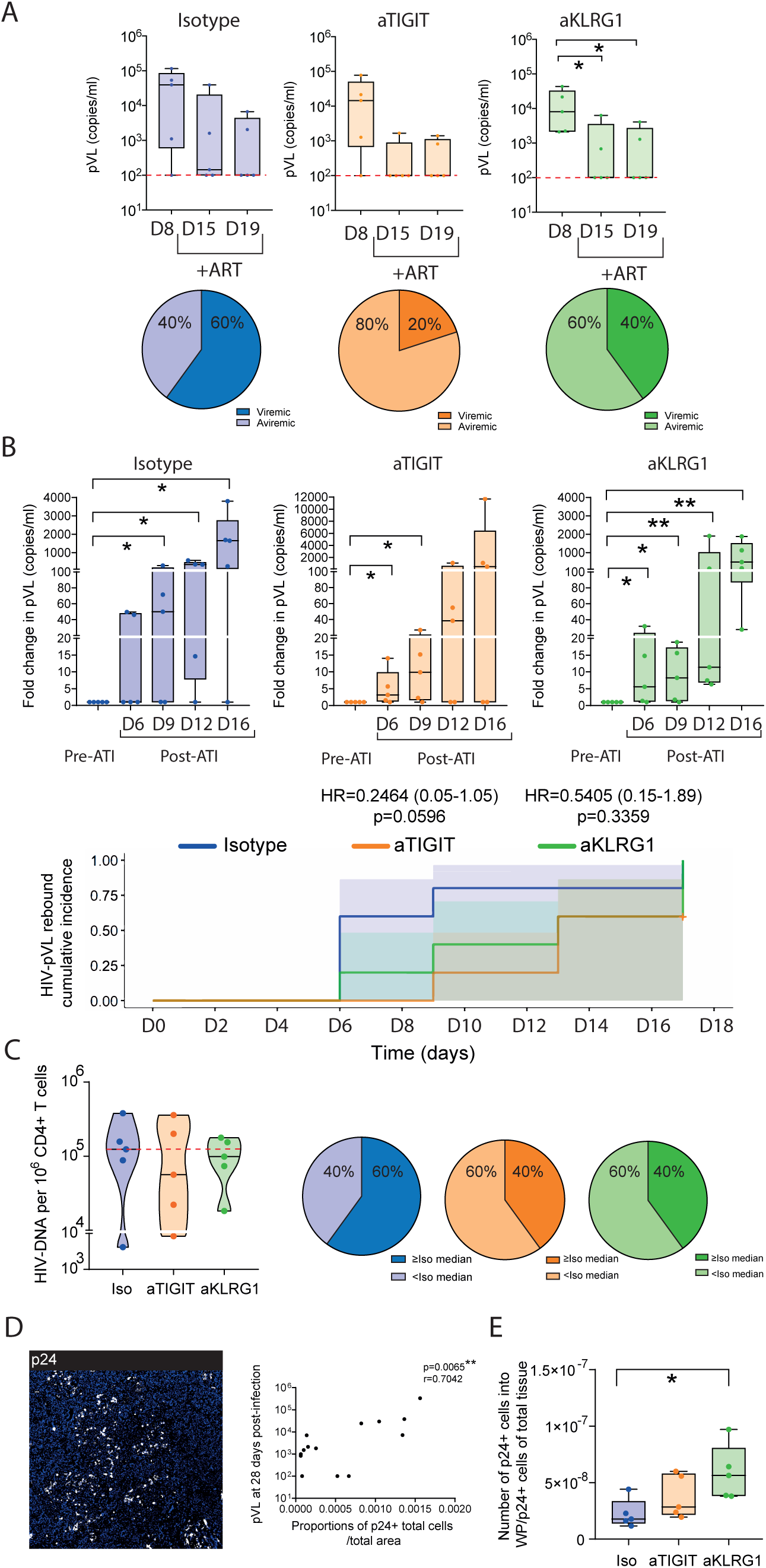
Analysis of viral suppression and HIV-1 viral load rebound after combination of ART with individual aTIGIT or aKLRG1 mAbs. (A): HIV plasma viral loads (pVL) prior to ART (8 days post-infection) and after ART combined with either isotype (blue, left plot), aTIGIT (orange, center plot) and aKLRG1 (green, right plot) mAbs (n=5 mice per group) at 15 and 19 days post-infection (top panels). Pie chart showing the percentage of viremic and aviremic BLT mice in each treatment group at day 15 (bottom panels). (B): Fold change in plasma HIV-1 RNA levels at 6, 9, 12 and 16 days post-ATI compared to pre-ATI levels in isotype (blue), aTIGIT (orange) and aKLRG1 (green) groups (top panels). Cumulative incidence curves for HIV-pVL rebound in each treatment antibody group is shown in the bottom panel. Shadowed areas represent 95% confidence intervals. (C): Levels of pro-viral HIV-1 DNA normalized per 10^6^ CD4+ T cells in PB from isotype (blue), aTIGIT (orange) and aKLRG1 (green) mAb-treated hBLT-mice groups at the end of experiment (left). Pie-charts showing the percentage of hBLT mice displaying integrated HIV-1 DNA values above or below the median values of the isotype hBLT group (red line) in the aTIGIT and aKLRG1 mAb treated animal groups (right). (D): Representative confocal microscopy image showing expression of p24 (white) distributed in the spleen from an infected hBLT mouse (left). Spearman correlation between the number of p24+ cells corrected to total tissue area (cells/μm^2^) and HIV-pVL at 28 days post-infection in the different treatment antibody groups (n=15) (right). (E): Ratio between number of p24+ cells into white pulp areas and p24+ cells in total tissue from isotype, aTIGIT and aKLRG1 mice groups. In panel A, B, C and E data are presented as Box and Whiskers plots showing median values, with minimum and maximum error bars. Pie chart data in panels A and C represent group proportions. Statistical significance was calculated using a using a Mann-Whitney test, a Chi-square test or a Regression Cox curve. *p<0.05; **p<0.01.

When we analyzed HIV-1 pVL after ATI, a delay in viral rebound was observed in hBLT mice treated with aTIGIT and aKLRG1 mAb immunotherapy when comparing to the maximal pVL suppression achieved before ATI on each group of animals (Figure 2B, upper plots). This beneficial effect was more evident in median at six and nine days post-ATI in these groups of mice. Rebound cumulative incidence in the aTIGIT Ab-treated group was especially lower compared to isotype group as illustrated in a Cox regression analysis (Figure 2B, bottom). Indeed, even at 16 days post-ATI, animals treated with the aKLRG1 mAb showed close to a log lower of pVL (Supplemental Figure 4D).

To further assess the level of viral control in these animals, analysis of pro-viral HIV-1 DNA was also carried out in CD4+ T cells from peripheral blood (PB) and spleen (SP) of mice from different treatment groups at the end of experiment. While total HIV-1 provirus levels were comparable in CD4+ T cells from the spleen between different treatment groups (Supplemental Figure 4D), 60% of mice from the aTIGIT and aKLRG1 mAb treated groups exhibited lower levels of HIV-DNA in circulating CD4+ T cells compared with median values present in mice treated with isotypic mAb, despite of high levels of HIV-1 pVL after viral rebound (Figure 2C). In contrast, in animals receiving the bispecific aTIGIT/aKLRG1 mAb treatment, loss of control was detected earlier after ATI than in the mice treated with aTIGIT and aKLRG1 Ab treatments, which lost control at day nine post-ATI (Supplemental Figure 4D, E, F). Consistently, levels of total HIV-1 proviral DNA detected in CD4+ T cells from PB were higher in 80% of mice treated with bispecific mAb compared to mice treated with isotypic mAb (Supplemental Figure 4G). The same tendency was observed in SP from mice treated with bispecific Ab (Supplemental Figure 4H). We then asked whether proportions and distribution of HIV-1-infected cells in lymphoid tissue could be associated with differences in pVL observed. To this end, we analyzed the expression of the p24 marker in tissue sections from the different animal groups by confocal microscopy (Figure 2D, Supplemental 4I). Interestingly, levels of HIV-1 pVL during rebound after ATI were significantly positively correlated with higher numbers of p24+ HIV-1-infected cells in lymphoid tissue (Figure 2D). The distribution of p24+ cells within the spleen tissue evidenced different patterns in the different hBLT mice groups. While no significant differences were detected in the total number of p24+ cells in these tissues (Supplemental Figure 4J), hBLT mice treated with aTIGIT mAb tended to exhibit less infected cells both inside and outside the white pulp areas of the spleen compared to animals receiving isotypic control, aKLRG1 or bispecific mAbs (Supplemental Figure 4K, L). This trend was also observed in proportion of mice treated with aTIGIT mAb whose tissue p24 detection was below median values of the control group in a higher proportion of animals (Supplemental Figure 4K, L). Notably, the group of hBLT mice treated with aKLRG1 mAb showed a significant accumulation of infected cells in white pulp areas relative to outlier regions (Figure 2E). Interestingly, the distribution of p24+ cells tended to also follow a similar pattern in the bispecific group, with a higher detection of infected cells in the white pulp areas, although no significant differences were observed (Supplemental Figure 4M). In conclusion, these findings suggest that combination of ART with individual aTIGIT or aKLRG1 mAb immunotherapy enhances viral suppression and induces partial control of HIV-1 replication after ATI in the hBLT mouse model, but potentially through different mechanisms.

### aTIGIT mAb immunotherapy is associated to cytotoxic adaptive-NKG2C+ NK cells in the spleen of HIV-1 infected hBLT mice

Next, we aimed at elucidating the immunological mechanisms potentially involved in control of viral replication in each of the hBLT mice group treated with the individual aTIGIT and aKLRG1 mAbs. Treatment with the aTIGIT mAb seemed to be associated with an enrichment of a precursor of adaptive NKG2C+ NK cells characterized by a NKG2C+ CD57– phenotype (Figure 3A, Supplemental Figure 5A, B, C, D). In contrast, the group of hBLT mice treated with the aKLRG1 mAb showed a lower proportion of NKG2C+ CD57-NK cells (Figure 3A, Supplemental Figure 5A, B, C, D). Similar tendencies to aKLRG1 mAb treated group were found in hBLT mice treated with bispecific aTIGIT/aKLRG1 mAb, which showed a decreased of NKG2C+ CD57- and an enrichment of NKG2C-CD57+ NK cells despite no obvious differences in total CD56+ NK cells proportions in tissue (Supplemental Figure 5E, F, G). Focusing on the functional profiles of these NKG2C+ NK cells, we analyzed cytotoxicity and activation markers such as CD107a, IFNγ and TNFα in these cells (Supplemental Figure 5H). As shown in Figure 3B, proportions of total CD107a+ cells and CD107a+ IFNγ+ cells within the adaptive NKG2C+ CD57– adaptive NK cell precursors considering the spleens of mice treated with the aTIGIT and isotypic mAbs significantly correlated with lower HIV-1 pVL after ATI. In the case of hBLT mice treated with the aKLRG1 and isotypic mAbs, we did not observe any significant correlations between pVL and these cytotoxic adaptive NKG2C+ CD57-NK cell precursors in the spleen (Supplemental Figure 5I, J). However, the proportions of CD56+ CD16+ NK cells expressing CD107a and TNFα correlated only with the proportions of cytotoxic NKG2C– NK cells when considering the mice groups treated with aKLRG1 and isotypic mAbs (Supplemental Figure 5K).

**Figure 3.**
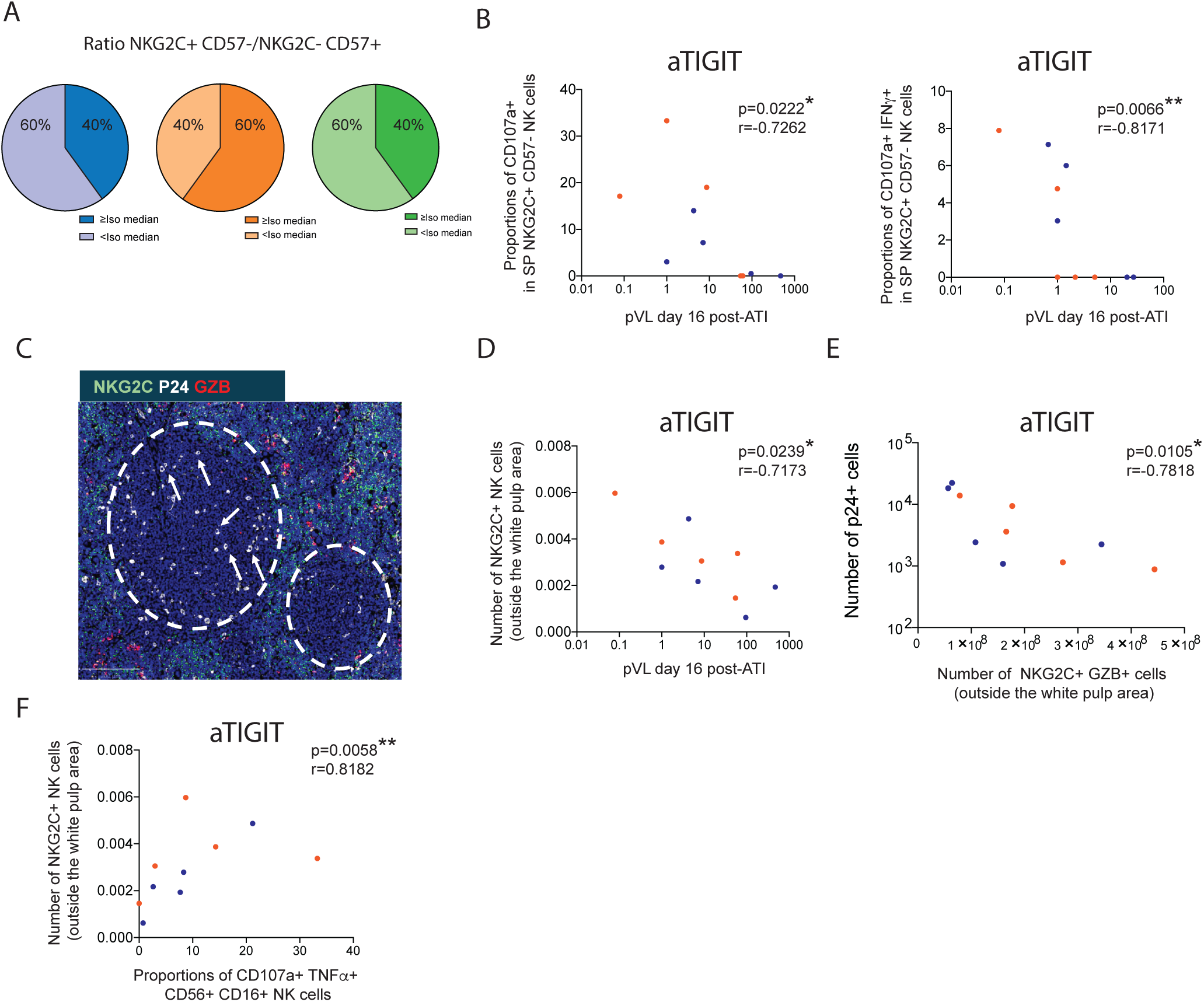
Functional characterization of adaptive NKG2C+ NK cells in the spleen of HIV-1 infected hBLT mice treated with aTIGIT mAb. (A): Pie-charts showing the percentage of hBLT mice with ratio between NKG2C+ CD57- and NKG2C-CD57+ NK cells above or below the median of the isotype group (blue) in the aTIGIT (orange) and aKLRG1 (green) BLT mice groups (see supplemental Figure 5B). (B): Spearman correlations between proportions of degranulation marker CD107a (left) and CD107a+ IFNγ+ (right) in NKG2C+ CD57-NK cells versus HIV-1 pVL post-ATI. Dots from mice treated with aTIGIT and isotype mAbs are colored in orange and blue, respectively. (C): Representative confocal microscopy image showing expression of NKG2C (green), p24 (white) and granzyme B (red) in the spleen from a representative aTIGIT mAb treated hBLT mouse. White pulp areas are highlighted in white circles. (D): Spearman correlation between number of NKG2C+ cells outside the white pulp areas versus HIV-1 pVL 16 days post-ATI in the aTIGIT (orange) and isotype (blue) mAb treated-hBLT mice group. (E): Spearman correlation between number of NKG2C+ cells expressing granzyme B outside the white pulp areas versus number of p24+ cells in total tissue area (cells/μm^2^) in the aTIGIT (orange) and isotype (blue) treated-BLT mice group. (F): Spearman correlation between number of NKG2C+ cells outside the white pulp areas versus proportions of CD107a+ TNFα+ CD56+ CD16+ NK cells in the aTIGIT (orange) and isotype (blue) mAb treated-BLT mice group. In panel A data are presented as Pie chart. Statistical significance was calculated using a Chi-square test or a Spearman correlation test. *p<0.05; **p<0.01.

Subsequently, after confirming by FACS that NK cells were the cell lineage preferentially expressing NKG2C compared to T cells in hBLT mice (Supplemental Figure 5L), we performed a histological analysis of lymphoid tissue to locate adaptive NKG2C+ cells and the cytotoxic marker Granzyme B (GZB) (Figure 3C; Supplemental Figure 5M). Considering the distribution of NKG2C+ cells in the tissue of aTIGIT mAb-treated hBLT mice, we observed that a higher number of these cells outside the white pulp areas correlated with lower viral loads following ART interruption (Figure 3D). These cells seemed to be evenly distributed in red pulp areas in the aTIGIT treated group, while a more polarized phenotype of these cells was found in these cortical areas in the isotype animal group (Supplemental Figure 5M). Moreover, the co-expression of NKG2C and GZB correlated with a reduced number of p24+ infected cells in the lymphoid tissue considering hBLT mice treated with aTIGIT and isotypic mAbs (Figure 3E). Additionally, we observed that the presence of these NKG2C+ cells outside the white pulp also correlated with a population of NK cells in these mice expressing CD107a and TNFα in the spleen in these mice (Figure 3F).

Together, these data indicate that control of HIV-1 replication mediated by blockade of TIGIT but not KLRG1 is associated with an induction of cytotoxic NKG2C+ adaptive NK cells in tissue from infected hBLT mice.

### Increased cytotoxic and IFNγ producing CD8+ T cells in the spleen white pulp of hBLT mice treated with aKLRG1 mAb

Next, we focused on investigating functional activation profiles of CD8+ T lymphocytes in both hBLT groups treated with aTIGIT and aKLRG1 mAbs. Interestingly, CD8+ T cells expressing IFNγ and TNFα, but lacking the degranulation marker CD107a, were enriched in the spleen of aTIGIT mAb-treated hBLT mice compared to the group of animals treated with the control isotypic mAb (Supplemental Figure 6A). In contrast, this same CD8+ T cell subpopulation presented lower frequencies in both the aKLRG1 and the bispecific-mAb-treated groups compared to the aTIGIT mAb-treated group (Supplemental Figure 6A). While no correlation of this population with pVL was found in the aTIGIT mAb -treated group (Supplemental Figure 6B), we did find a negative correlation between these cells and cytotoxic NK cells (Supplemental Figure 6C), and particularly with CD107a+ IFNγ+ non-adaptive NKG2C-NK cells (Supplemental Figure 6D, right) and to some extent with the same population included in adaptive NKG2C+ NK cells previously observed in the aTIGIT mAb-treated mice group (Supplemental Figure 6D, left). These data suggested a preferential negative correlation between IFNγ+ TNFα+ CD107a-CD8+ T cell with non-adaptive NK cells.

Focusing on identifying an immune control mechanism specifically induced in the aKLRG1-mAb versus the isotype mAbs-treated groups, we found that the proportions of cytotoxic CD107a+ CD8+ T lymphocytes present in peripheral blood (PB) and spleen (SP) showed a significant negative correlation with HIV-1 pVL post-ATI and so were associated to viral control in this group (Figure 4A). This population of cytotoxic CD8+ T cells tended to positively correlate with the proportions of CD56+ CD16+ NK cells co-expressing CD107a and TNFα in the lymphoid tissue of mice treated with the aKLRG1 mAb (Supplemental Figure 6E). Additionally, a population of CD8+ CD107a+ IFNγ+ TNFα– T cells correlated negatively with levels of HIV-1 pVL post-ATI (Figure 4B) and positively with the proportions of CD8+ T cells expressing only IFNγ in mice treated with aKLRG1 (Figure 4C). It is also noteworthy that Granzyme B expression in CD8+ T cells strongly correlated with lower levels of post-ATI viral load in this mice group (Supplemental Figure 6F).

**Figure 4.**
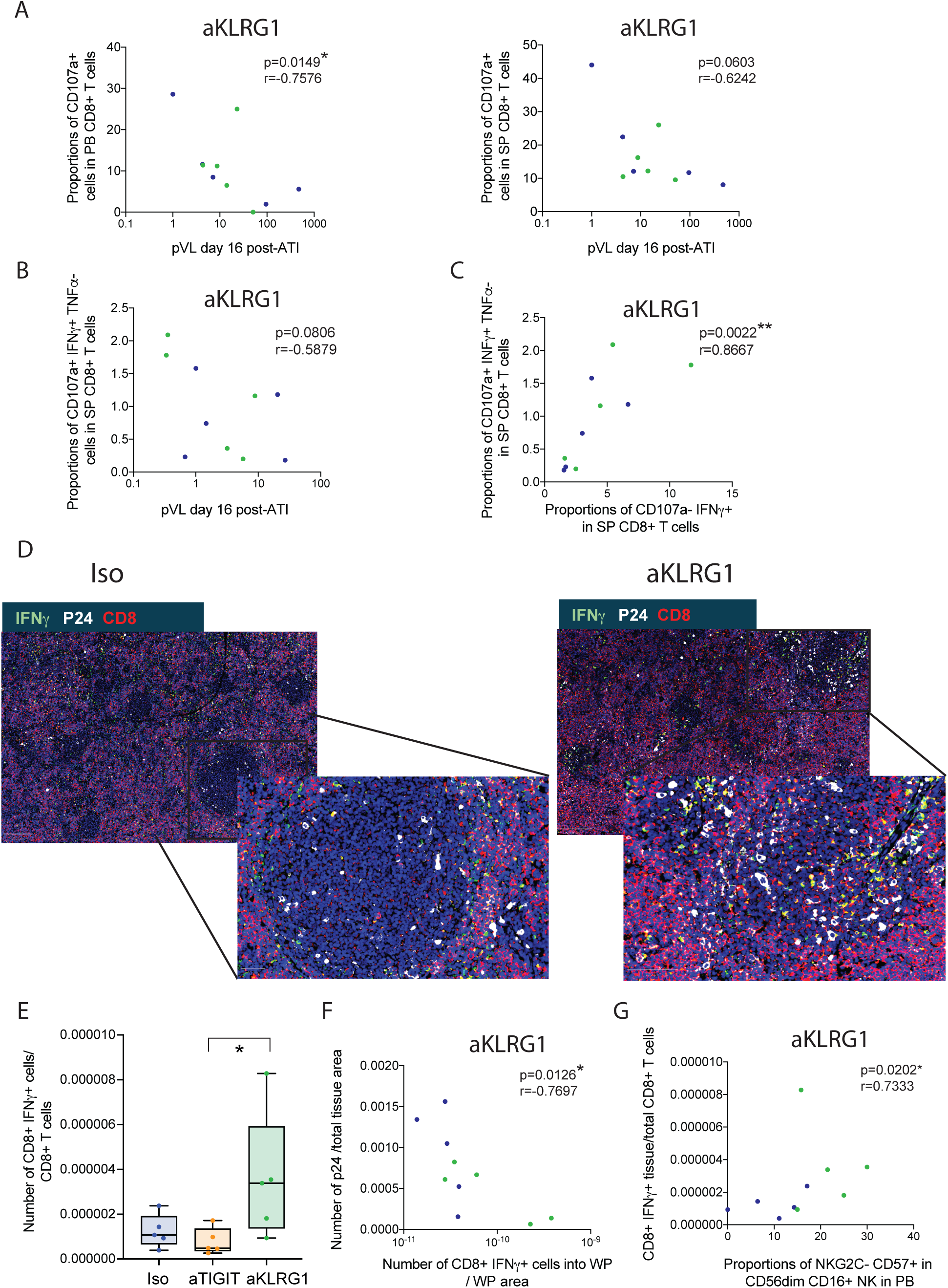
Differential functional patterns of CD8+ T cells in HIV-BLT mice treated with aKLRG1 mAb. (A): Spearman correlations between proportions of degranulation marker CD107a in CD8+ T cells in peripheral blood (PB; left) or spleen (SP; right) and HIV-1 pVL post-ATI in aKLRG1 mAb treated hBLT mice. (B): Spearman correlation between proportions of CD107a+ IFNγ+ TNFα-CD8+ T cells in SP and HIV-1 pVL post-ATI. Dots from mice treated with aKLRG1 and isotype mAbs are colored in green and blue, respectively (C): Spearman correlation between proportions of CD107a+ IFNγ+ TNFα-CD8+ T cells and proportions of CD107a-IFNγ+ CD8+ T cells located in SP of hBLT mice treated with aKLRG1 mAbs. Dots from mice treated with aKLRG1 and isotype mAbs are colored in green and blue, respectively (D): Representative confocal microscopy image showing expression of IFNγ (green), p24 (white) and CD8 (red) in the spleen from representative hBLT mice treated with an Isotypic IgG (left) and aKLRG1 (right) mAbs. (E): Number of CD8+ T cells expressing IFNγ normalized to total number of CD8+ T cells in SP from the different hBLT treatment groups (F, G): Spearman correlation between number of CD8+ IFNγ+ T cells into the white pulp areas versus number of infected p24+ cells (G) or the proportions of NKG2C-CD57+ in PB (G) at the end of experiment in the hBLT mice group treated with aKLRG1 mAb. Dots from mice treated with aKLRG1 and isotype mAbs are colored in green and blue, respectively. In panel E data are presented as Box and Whiskers plots showing median values, with minimum and maximum error bars. Statistical significance was calculated using a Spearman correlation test or a Mann-Whitney test. *p<0.05; **p<0.01.

We next quantified the distribution of CD8+ T lymphocytes in lymphoid tissue and the expression of IFNγ by histological analysis in the SP (Figure 4D; Supplemental Figure 6G). Although we did not detect significant differences in the number of CD8+ T cells between groups (Supplemental Figure 6H), a significantly higher proportion of IFNγ-expressing CD8+ T cells was observed in the aKLRG1 mAb-treated hBLT mice compared with the aTIGIT mAb-treated animal group and almost significant compared with isotype group (Figure 4D, E; Supplemental Figure 6G). Notably, in the aKLRG1mAb-treated mice, IFNγ+ CD8+ T cells were enriched within the white pulp of the spleen compared with the Isotype and aTIGIT mAb-treated mice groups (Figure 4D; Supplemental Figure 6G, H, I, J), which correlated with a lower number of p24+ HIV-infected cells in this tissue, suggesting that these cells may be mediating the previously observed restriction of infected cells to these areas (Figure 4F).

Moreover, the increased frequency of CD8+ T cells in the SP from aKLRG1 and isotype mAb-treated mice positively correlated with the proportion of NKG2C– CD57+ NK cells, previously shown to be enriched in this group (Figure 4G). Collectively, these data suggest that cytotoxic IFNγ+ CD107a+ CD8+ T cells play a more prominent role in mediating immune control of viral replication in the aKLRG1 mAb-treated group.

### Identification of early immune-biomarkers modified by ART and aTIGIT/aKLRG1 mAb associated to control of HIV-1 viral replication in aTIGIT and aKLRG1 mAb-treated hBLT mice

It is relevant identifying early immune biomarkers specifically associated with immune responses occurring in each hBLT mice group predicting responses observed in the presence of aTIGIT or aKLRG1 mAbs. In this regard, we analyzed the immune phenotype of these mice at earlier time points in which combined with ART was applied in combination with immunotherapy (IT) at day 15 post-infection. We observed that proportions of mature CD56dim CD16+ NK cells were significantly reduced in the PB of aKLRG1 mAb-treated mice compared with aTIGIT and isotype mAb-treated animal groups, in which the proportions of these cells were maintained (Figure 5A). At this time point, the circulating NK cells present in the aKLRG1 mAb treated group were characterized by a more effector phenotype NKG2C- and lower proportions of adaptive NKG2C+ NK cells, as opposed to aTIGIT mAb treated hBLT mice, which showed higher proportions of NKG2C+ NK cells (Figure 5B-C). In accord, the enrichment of NKG2C-CD57+ NK effector subset was associated to higher number of p24+ infected cells in the tissue from aKLRG1 vs isotype mAb-treated hBLT mice (Figure 5D), indicating that enrichment of this NK population associates to a worse HIV-1 control. Moreover, in the group of hBLT mice treated with aTIGIT vs isopype mAb we also found a negative association between the proportions of NKG2C+ CD57-NK cells and NKG2C-CD57+ NK cells, supporting the enrichment of adaptive NKG2C+ cells in this group (Figure 5E). In this regard, proportions of CD56dim CD16+ NK cells were also depleted in the blood after bispecific mAb treatment at this early time point (Supplemental Figure 7A). Moreover, proportions of NKG2C+ were depleted in this group in contrast to NKG2C-, which were enriched following the same tendency than aKLRG1 mice treated group (Supplemental Figure 7B, C). Thus, depletion of circulating CD56dim CD16+ NK cells and enrichment in adaptive NKG2C+ NK cells are earlier biomarkers of immune responses present in aKLRG1 and aTIGIT mAb treated groups, respectively. Regarding the analysis of CD8+ T lymphocytes during combination of ART plus individual aTIGIT or KLRG1 mAb immunotherapy, a population of CD8⁺ T cells expressing CD57 tended to decrease in the PB from mice treated with aKLRG1 mAb compared with animals receiving the control isotype mAb (Figure 5F). Noteworthy, proportions of these CD57+ CD8+ T cells were negatively associated with the enrichment of the IFNγ+ CD8+ T cell subpopulation found in the white pulp (Figure 5G) of lymphoid tissue from the aKLRG1 mAb treated hBLT group, which had previously been linked to improved viral control. The proportions of CD8+ CD57+ T cells at early time points were associated to higher number of p24+ infected cells in lymphoid tissue at the end of experiment in aTIGIT mAb-treated hBLT mice (Supplemental Figure 7D). Together, different phenotypes in NK and CD8+ T cells during ART and aTIGIT or aKLRG1 mAb immunotherapy were associated to subsequent control of HIV replication in these hBLT mice groups.

**Figure 5.**
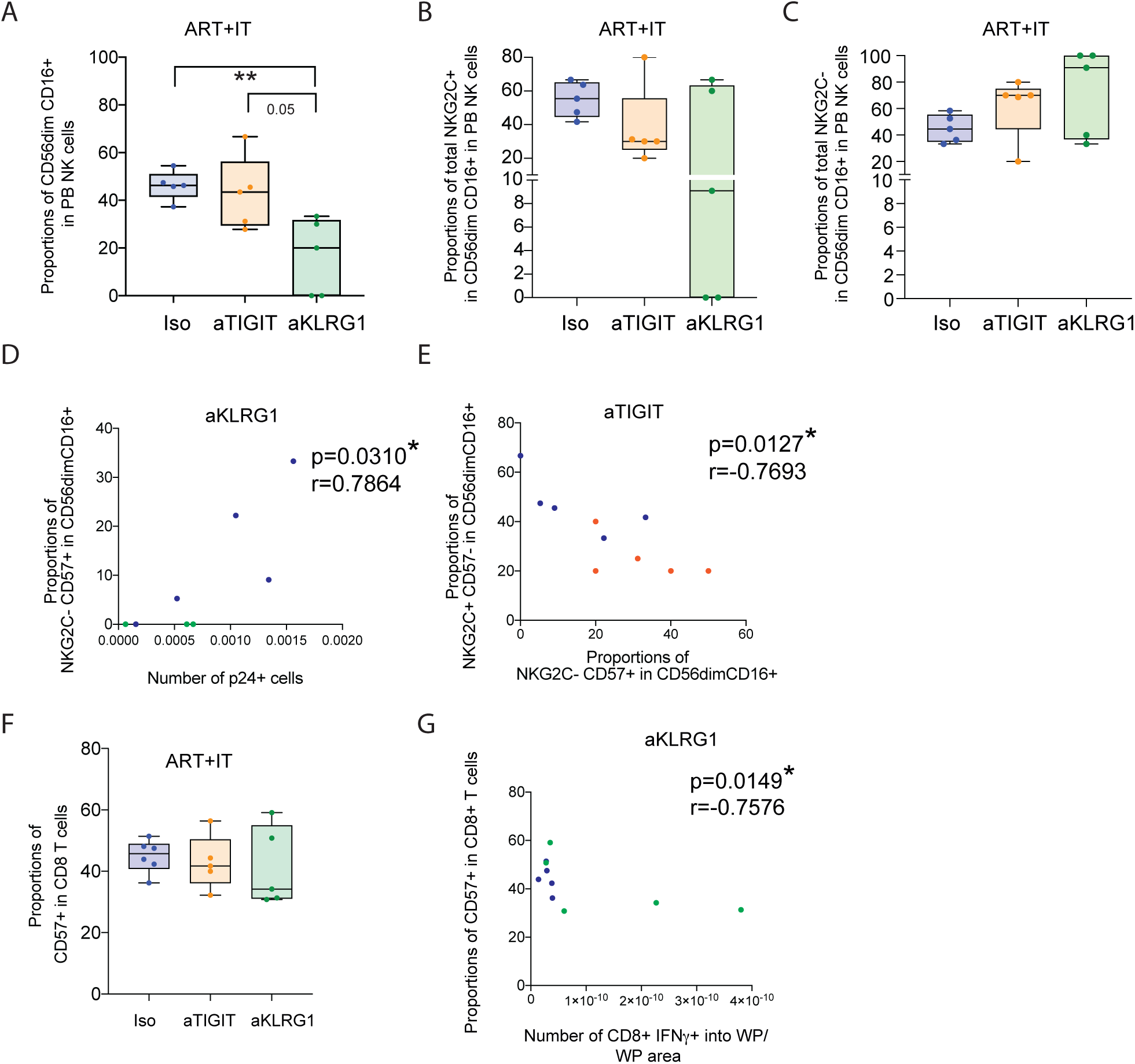
Phenotypical analysis of NK and CD8+ T cells in the different antibody treatment groups. (A): Proportions of CD56dim CD16+ NK cells in PB from each n=5 isotype (blue), aTIGIT (orange) and aKLRG1 (green) BLT-mice treatment groups at first week after ART plus IT initiation. (B, C): Proportions of NKG2C+ (B) and NKG2C-(C) in CD56dim CD16+ NK cells in PB from each n=5 isotype (blue), aTIGIT (orange) and aKLRG1 (green) BLT-mice treatment groups at first week after ART plus IT initiation. (D): Spearman correlation between proportions of NKG2C-CD57+ CD56dim CD16+ NK cells in PB at ART initiation and number of p24+ infected cell in SP from aKLRG1 (green) and isotype (blue) mAb-treated hBLT mice at the end of experiment. (E): Spearman correlation between proportions of NKG2C+ CD57- and NKG2C-CD57+ CD56dim CD16+ NK cells in PB from aTIGIT (orange) and isotype (blue) mAb treated hBLT groups at ART plus IT initiation. (F): Proportions of CD57+ CD8+ T cells in SP from each n=5 isotype (blue), aTIGIT (orange) and aKLRG1 (green) BLT-mice treatment groups at first week after ART plus IT initiation. (G): Spearman correlation between proportions of CD8+ CD57+ T cells in PB at ART plus IT initiation versus number of CD8+ IFNγ+ T cells into WP in SP at the end of experiment in the aKLRG1 (green) and isotype (blue) mAb-treated-hBLT mice group. In panel A, B, C and F data are presented as Box and Whiskers plots showing median values, with minimum and maximum error bars. Statistical significance was calculated using a Mann-Whitney test. *p<0.05; **p<0.01.

### Induction of apoptosis of NK stimulated by bispecific aTIGIT/aKLRG1 mAb

Due to the observed decrease in NK cell proportions in the presence of aKLRG1 and the bispecific mAbs treatment *in vivo*, as well as the unexpected inefficacy of the bispecific mAb activity both in combination with ART and after ATI, we assessed the mechanisms underlying these responses. To this end, we analyzed the Annexin V content, a marker of apoptosis, after treating PBMCs from PWH with each specific antibody at the selected concentration. An enrichment in Annexin V+ proportions was found in both adaptive NKG2C+ and non-adaptive NKG2C+ NK cells following incubation with the bispecific antibody (Supplemental Figure 8A, left). In the case of NKG2C-NK cells, we also observed an increase in Annexin V+ cells after aKLRG1 mAb treatment, although lower than that induced by the bispecific Ab (Supplemental Figure 8A, right). In contrast, apoptosis levels were not affected in either adaptive or non-adaptive NK cells following treatment with aTIGIT mAb (Supplemental Figure 8A). However, in CD8+ T lymphocytes, we did not detect significant changes after incubation with any of the tested antibodies in apoptosis levels, although a trend towards lower levels in cells exposed to aKLRG1 and bispecific aTIGIT/aKLRG1 mAbs was observed (Supplemental Figure 8B). Together, these findings suggest that loss of control of immune control in hBLT treated with the bispecific mAb may be due to apoptosis of NK cells.

## METHODS

### Individual and bispecific anti-TIGIT, anti-KLRG1 antibodies

Humanized anti-TIGIT blocking mAb (Tiragolumab) previously used in clinical trials for lung cancer ^51^ was obtained through company Selleck Chemical and used for this study. Humanized anti-KLRG1 IgG1 was based on the patent application 20210347899 (ABC_HG1N07 sequence of an activating mAb) and manufactured by WuXi Biologic. The bispecific aTIGIT/aKLRG1 mAb used in this study was designed by Dr. María J. Buzón and was generated using the sequences from the same variable Fab regions from each of the mentioned individual aTIGIT and aKLRG1 mAb as well as a human IgG1 isotype heavy chain and manufactured at WuXi Biologics. A humanized IgG1 isotypic mAb (BioXCell) mAbs was used as control for in vitro and *in vivo* experiments. PBMCs obtained from n=8 PWH on ART, characterized by undetectable plasma viremia (<20 copies HIV-1 RNA/mL) and median of 626.185 CD4+ T cell counts (360-1149.23 Min-Max Range) were recruited from the Infectious Diseases Unit from Hospital Universitario La Princesa thanks to an approved informed consent (Protocol Registration Number 3518) and in accordance with the Helsinkís declaration and the department of Health and Human Services Belmont Report. PBMCs from HIV-1 negative donors (HD) were obtained from HIV-1 negative donors using Buffy coats donated by the Centro de Transfusiones Comunidad de Madrid. PBMC from PWH and HD were used to test Ab binding specificity and also for functional test of NK and CD8+ T cells (see sections below).

### Isolation of human NK, DC and CD8+ T cells subsets

Total DC were purified from total PBMC suspensions from HD by negative immunomagnetic methods (purity >90%) using the Human Myeloid DC Enrichment Kit (STEMCELL). Human T cells were isolated by negative immunomagnetic sorting from PBMC of HD (Untouched human T cell isolation Kit, Invitrogen) and co-cultured with allogeneic DC (see functional assay section). NK cells were isolated by negative immunomagnetic selection (STEM cell) from the peripheral blood of aviremic PWH on ART from our study cohort following the manufacture instructions, reaching >90% purity.

### Functional test of antibodies

To determine the functional effect and optimal working concentrations of the individual aTIGIT, aKLRG1 and bispecific aTIGIT/aKLRG1 mAbs used in subsequent experiments, NK cells isolated from PWH were co-cultured with the MHC-I deficient target cell line K562 transfected with GFP (NIH AIDS reagent program Ref 11699) at a target:NK ratio of 1:2 in the presence of serial dilutions of these antibodies for 16 hours. Following incubation, NK cell functional responses were assessed by intracellular staining of CD107a and IFN-γ and analyzed by flow cytometry (FACS). A dose-response curve was generated using antibody concentrations ranging from 0.000001 μg/mL to 10 μg/mL.

To evaluate the effect of the antibodies on CD8+ and CD4+ T cells, a mixed lymphocyte reaction (MLR) was performed using T cells from healthy donors (HDs) in the presence of allogeneic DC previously at ratio 1:4 DC-CD8+ T cells, stimulated for 16h in the presence of 1ug/mL Poly I:C (InvivoGen) and 2‘3’-c-di-AM(PS)2 (Invivogen) in the presence of 5 μg/mL of either isotypic control or anti-TIGIT, anti-KLRG1 or the bispecific aTIGIT/aKLRG1 mAbs. After 5 days, co-cultures were stimulated with 0.25μg/ml PMA and 0.25μg/ml Ionomycin CD8⁺ T cell activation was assessed by evaluating intracellular expression of CD107a, IFN-γ and TNF-α by FACS. A control condition consisting of T cells cultured in medium alone was included. An isotype-matched control mAb was included at a 5μg/mL concentration as a negative control.

### Infection of hBLT mice with HIV-1 and combination of ART with immunotherapy

NOD/SCID IL2R-y-/-(NSG) mice transplanted with human BM, fetal liver, and thymus (BLT-mouse) and fetal CD34+ hematopoietic stem cells (HSCs) were generated as previously described ^39^ at the Human Immune System Core from the Ragon Institute and Massachusetts General Hospital in collaboration with Dr. Vladimir Vrbanac and Dr. Alejandro Balazs. hBLT mice were housed in ventilated racks and fed autoclaved food and water at a pathogen-free facility. Intraperitoneal injections of 0.125μg/g rhIL-15 (R&D systems) were performed in hBLT mice for 2 weeks to increase reconstitution and maturation of human NK cells in these animals. Human immune reconstitution was monitored for 2 weeks prior to HIV-1 infection and optimal reconstitution was defined by proportions of human CD45+ lymphocytes over 25% in PBMC. Viral stocks were generated by transfection of HEK-293 (ATCC® CRL-1573) cells with the plasmid “HIV-1 JR-CSF Infectious Molecular Clone (pYK-JRCSF)” (National Institutes of Health [NIH] AIDS Reagent Program, catalog number 2708) using the X-tremeGENE 9 DNA Transfection Reagent (Roche) according to manufacturer’s instructions. 72 h after transfection, supernatants were collected, filtered, concentrated and resuspended in 1x PBS using Lenti-X (TaKaRa) according to manufacturer’s instructions. The JRCSF viral stock was stored at −80°C for further experiments. The virus TCID_50_ was determined in TZM-bl cells (NIH AIDS Reagent Program, catalog number 8129) by limiting dilution using the Reed and Muench method, as previously described. Dr. Julia García Pradós group at IRSICaixa, Barcelona, Spain produced and provided the viral stocks for the study. A total of n=20 reconstituted hBLT mice were intravenously infected with 5,000 TCID_50_ of JRCSF HIV-1 virus. *In vivo* experiments were conducted in a Biosafety Level 3 (BSL3) animal facility at the Centro de Biologia Molecular Severo Ochoa (CBM) in Madrid. After eight days post-infection HIV-1 plasma viral loads were determined and animals were randomized in four groups of n=5 animals, attending to reconstitution levels and weight. Subsequently, after 1 week infected hBLT mice initiated oral ART ((87.5 mg/kg/day emtricitabine (FTC), 131 mg/kg/day tenofovir (TDF), and 175 mg/kg/day raltegravir (RAL) and darunavir (DRV)) administered via drinking water, as described ^55^ and at the same time, received intraperitoneal injection of 150 μg of either aTIGIT, aKLRG1, aTIGIT/aKLRG1 bispecific and isotype control mAb immunotherapy. The immunotherapy regimen was maintained every 3 days from 8 to 32 days post-infection. After 7 days since ART+IT, blood extraction was performed for phenotypical analysis of NK and CD8+ T cells by FACS. At 36 days post-infection, mice were sacrificed and total blood and spleen were collected, 1/3 of spleens were used for histological analysis, 1/3 was used by FACS phenotypical analysis by FACS and 1/3 preserved in liquid nitrogen for proviral-DNA analysis. Total peripheral blood also was collected for immunophenotypical analysis and proviral-DNA quantification (100μl). Body weight, HIV-1 pVL, were weekly monitored during the duration of the whole experiment (See Supplemental Figure 6D). For *in vivo* experiments, mice were housed at the BSL3 animal facility from Centro de Biología Molecular Severo Ochoa (CBM) in accordance with the institution’s animal care standards. Animal experiments were reviewed and approved by the local ethics committee in agreement with the EU Directive 86/609/EEC, Recommendation 2007/526/EC and Real Decreto 53/2013.

### Viral load determination

Plasma viral load (pVL) was measured from plasma samples collected from HIV-1 hBLT mice during ART and after ART interruption until the end of experiment weekly. HIV-RNA was extracted using QIAamp Viral RNA Mini kit (QIAGEN) following the manufacturer’s instructions. Reverse transcription of RNA to cDNA was performed with Promega kit in accordance with the instructions provided by the manufacturer, and cDNA was quantified by quantitative PCR (qPCR) using primers and probes specific for the Gag region (Gag forward 5′- TCAGCCCAGAAGTAATACCCATGT-3′ and Gag reverse 5′- CACTGTGTTTAGCATGGTGTTT-3′; Gag probe 5′-FAM-ATTATCAGAAGGAGCCACCCCACAAGA-TAMRA-3′), and Fast Advanced Master Mix (Invitrogen). Samples were analyzed in an Applied Biosystems QuantStudio5 system. Quantification of DNA was performed using a standard curve. Samples were analyzed in an Applied Biosystems QuantStudio5 system. Quantification of DNA was performed using a standard curve, and values were normalized to 1 million CD4+ T cells.

### Proviral DNA analysis

Total HIV DNA in the cell lysates was quantified by quantitative PCR (qPCR) using primers and a probe specific for the 1-LTR region of HIV (LTR forward: 5′- TTAAGCCTCAATAAAGCTTGCC-3′, LTR reverse: 5′-GTTCGGGCGCCACTGCTAG-3′, and probe: 5′/56-FAM/CCAGAGTCA/ZEN/CACAACAGACGGGCA/31ABkFQ/3′). Reactions were carried out using TaqMan™ Gene Expression Master Mix (Thermo Fisher). The CCR5 gene was used as a reference for cell input normalization. DNA quantification was performed using a standard curve, and values were normalized to one million CD4+ T cells, taking into account the percentage of CD4+ T cells obtained by flow cytometry from the same sample. Samples were analyzed using an Applied Biosystems QuantStudio 5 system.

### Flow cytometry staining

Different antibody cocktails containing mAbs identifying NK and CD3+ T cells from PWH PBMCs were tested by FACS using a cell viability marker as Viability Dye (Ghost DyeTM, Invitrogen) incubated in PBS for 10 min at 4°C in ice. PBMCs were stained with superficial antibodies, anti-CD56, anti-CD16, anti-CD3 and anti-human IgG antibodies (BioLegend) in Running Buffer (RB; Miltenyi Biotec; containing BSA, EDTA and 0.09% azide) for 15 min at 4°C in ice. After incubation, cells were washed and subsequently, cells were stained with an FITC-conjugated anti-human IgG secondary antibody (1:200 dilution; BioLegend), directed to the heavy chains of primary antibodies, for 20 min at 4°C in ice in RB. Finally, cells were fixed with 4% paraformaldehyde solution (PFA) and acquired in a LSR Fortessa flow cytometer (Becton Dickinson). The binding of anti-human IgG was detected through APC signal and gated on NK and CD3+ populations using FlowJo v10 software.

The ability of NK and CD8+T cells from PWH and HD to secrete cytokines and degranulate was performed using Ghost Dye Red 780 dye, anti-CD56, anti-CD16, anti-CD3, anti-CD8, in combination with either intracellular IFNγ, CD107a, TNFα mAbs in presence of 0.25 μg/mL Brefeldin A and 0.0025 mM Monensin (SIGMA) after 16h. Cells were fixed with Fixation buffer (BDBiosciences) for 30 min at 4°C in ice and after washing cells with permeabilization solution (BDBiosciences) during 5 min, intracellular antibodies were added to the permeabilization solution in presence of cells for 15 min at 4°C in ice. Cell line K562 was detected with GFP which is signaled in FITC in the case of NK cell culture. In humanized BLT mouse experiments, a combination of anti-human CD45, CD3, CD8, CD56, CD16, NKG2C and CD57 mAbs were used to analyze peripheral blood and spleen samples from transplanted mice following the experiment. At the end of experiment, 1/3 of the spleen from HIV-1-infected hBLT mice in each group, were excised, minced into small pieces and processed through a 70 μm cell strainer (Falcon). Both minced spleens and whole PB were employed for membrane and intracellular staining for flow cytometry analysis, following the procedure described above using antibodies, anti-human CD45, CD3, CD8, CD56, CD16, NKG2C, CD57, CD107a, IFNγ, TNFα and granzyme B (see Supplemental Table 1). Samples were analyzed in a BD FACS Canto II and Lyric instruments. Analysis of flow cytometry data was performed using FlowJo v10.6 software (Tree Star).

### Histological analysis of tissue sections from humanized mice

Spleen pieces were collected from hBLT mice and embedded in paraffin and sectioned in slides of 2 μm of thickness in a Leica microtome. Tissue sections were deparaffinized, hydrated, and target retrieval were performed with a PT-LINK (Dako) previous to staining. For paraffin-preserved tissue, rat anti-human Granzyme B (Thermo Fisher), mouse anti-HIV-1 p24 (Abcam), rabbit anti-human NKG2C (Abcam), rat anti-human CD8 (Bio-Rad) and rabbit anti-human IFNγ (Abcam) were used as primary antibodies; and goat or donkey anti-rabbit AF488 (Invitrogen), donkey anti-rat AF594 (Jackson ImmunoResearch), and donkey anti-mouse AF647 (ThermoFisher) as secondary antibodies (see Supplemental Table 1). Images were taken with a Leica TCS SP5 confocal and processed with the LAS AF software. Granzyme B+, NKG2C+, CD8+, IFNγ+ and HIV-1 p24+ cell counts and colocalization and distribution of some of these markers were analyzed with QuPath software. White pulp areas were defined by morphology and cell nuclear density using QuPath software.

### Statistical analysis

Statistical significance of differences between different or within the same animal groups receiving different treatments were assessed using a two tailed non-parametric Mann-Whitney U or Wilcoxon matched-pairs signed-rank tests, respectively. In some case, a Chi-square test was applied to determine differences in proportions of animals included on each group. For each individual mouse, HIV-1 pVL rebound was defined when pVL levels relative to day 19 post-infection (day 0 post-ATI) first exceeded a cutoff value of 15.1527. This was selected as the value which best separated rebound cumulative incidence curves among different treatment groups, in terms of the lowest p-value from log-rank test. To compare the kinetics of HIV-1 pVL rebound among different treatment groups, a Cox proportional-hazard model was used, computing hazard ratios (HR) and associated 95% confidence intervals in R v4.3.3 (*survival* package v3.6-4). For HR estimation, mice receiving isotype control mAb were considered as the reference group. Mice not experiencing HIV-1 rebound were censored at the time of their last follow-up. Non-parametric Spearman correlations were used to test both individual correlations and to generate a correlation network. Statistical significance was considered when p value<0.5. Statistical analyses were performed using GraphPad Prism 9.0 software. No blinding was applied to our data analysis.

## DISCUSSION

In this study, we used a hBLT mouse model to test the benefit of combining ART with an immunotherapy directed to either TIGIT or KLRG1 or both receptors to promote control of HIV-1 replication. Targeting inhibitory immune checkpoint with blocking antibodies has extensively studied in oncology and, for instance PD-1 blockade ^56^ is currently widely used in lung cancer. Indeed, in previous studies, the expression of PD-1 expression on NK cells did not discriminate between PWH groups with effective or dysfunctional response to DC stimulation, in contrast to TIGIT ^28^. More recently, TIGIT blockade has also been explored in this context ^51,57^ and the combination of blocking TIGIT during DC stimulation, has been shown to enhance NK and CD8+ T cell function against HIV-1 infected cells *in vitro* ^18,28^. However, the impact of targeting TIGIT had only been tested *in vivo* in a primate model of SIV infection and this immunotherapy alone not was sufficient to improve CD8+ T cell function against infected cells ^58^, but not in a humanized mouse model and in combination with other immunotherapies.

The low efficiency of NK cell reconstitution in the hBLT model requires additional approaches such as the exogenous intravenous IL-15 administration to achieve higher numbers and maturation of these cells ^28,44^. However, injection of rhIL-15 may at the same time, also impact CD8+ T cell activation and the induction and proliferation of tissue resident cells ^59,60^ and the expansion of NKG2C+ NK cells, previously described in NHP models ^61^, which may affect the antiviral immune response in these animals upon HIV-1 infection. In addition, administration of rhIL-15 could also contribute to CD4+ T cell polarization, which could potentially influence subsequent to latent reservoir reactivation, as suggested by some studies ^62,63^. Although rhIL-15 only was administrated only for two weeks prior to infection, its effects were limited to that period, and the specific changes observed in each independent treatment group were not associated with the rhIL-15 presence. However, their potential affecting other immune subpopulations should be investigated in more depth.

On the other hand, in our study we observed a synergistic effect of the combination of ART with individual aTIGIT or aKLRG1 mAb immunotherapy increasing the number of aviremic mice at early time points, potentially indicating an effective antiviral control likely mediated by reinvigorated exhausted CD8+ T and NK cells in presence of antiviral drugs, which could explain the earlier decay of HIV-1 plasma viral loads. This interpretation is compatible with results from previous studies that suggest a beneficial effect of immunotherapies combined with early ART initiation ^25,35,64,65^. In our study, hBLT mice were maintained on ART for 2 weeks, which was sufficient to reach undetectable viral loads even after just one week of treatment in the majority of animals. However, we did not analyze the evolution of persistent reservoirs in this model that may require longer ART duration as previously described ^25^. Although the establishment of viral reservoirs was not assessed in different time points along the experiment, previous studies in humans have shown that the HIV-1 reservoir is established within the first few days of infection ^66,67^. Moreover, previous studies determined that CD8+ T cells do not impact SIV reservoir establishment under ART ^68^, this may be modulated by NK cell function, which could explain the greater effect of aTIGIT treatment at this early time point (15 days post-infection). However, to properly study viral latency, future studies should determine viral sequence evolution in suppressed hBLT mice during ART. An additional potential limitation is the route of HIV-1 infection used for our study since it may influence the dynamics of viral dissemination, immune activation and reservoir establishment in the hBLT mouse model ^69^. In this regard, to avoid intrinsic infection rate variability in our study we used an intravenous (IV) route rather than an intravaginal (IVAG) infection method which may better recapitulate the physiology of sexual transmission. In contrast, IV infection with a high 5,000 TCID_50_ dose results in a faster and more uniform systemic viremia, but may overestimate the efficiency of viral dissemination, and could have potentially diminished the level of immune control induced by immunotherapy. In this regard, the level of viral rebound beyond pre-ART viral load level was reached after 31-35 days post-infection. Therefore, additional experiments using intravaginal challenge in hBLT should be conducted to confirm the presented results.

Regarding immune responses associated with viral control induced by immunotherapy, our results suggest differential NK and CD8+ T cell mediated mechanisms elicited by selective targeting of TIGIT and KLRG1 *in vivo*. On one hand, our study points towards an increased cytotoxic activity of NKG2C+ adaptive NK cells after TIGIT blockade, which fits our previous results *in vitro* and in humanized immune deficient NSG mice model of xenotransplant with NK and CD4+ T cells from PWH ^28^. We now provide evidence of the functional impact of TIGIT blockade in adaptive NK cells in the hBLT mouse model of HIV-1 infection which contains different immune cell lineages such as T cells that also express this molecule and had not been previously characterized in the context of chronic infection and ART combination. In this model, we investigated different immune mechanisms that may be involved in the control of HIV-1 infection.

Notably, the use of the agonistic anti-KLRG1 mAb was primarily associated with infection control mediated by cytotoxic CD8+ T cells. This suggests that aKLRG1 may enhance the function of virus-specific CD8+ T cells as has previously been reported, expressed in circulating virus-specific CD8⁺ T cells during LCMV infection ^34^. In addition, KLRG1 and is also associated with immunosenescence, being expressed on both effector terminal differentiated CD8+ T cells and NK cells in PWH ^70,71^. KLRG1 is an inhibitory receptor that binds to members of the cadherin family, and its engagement has been linked to functional impairment of NK and CD8+ T cells ^72,73^. Therefore, treatment of hBLT mice with the aKLRG1 mAb may block this inhibitory signaling, potentially restoring the effector function of these cells. By the other hand, a study has described that circulating KLRG1+ long-lived effector memory T cells may became in tissue resident T cells ^71^, which could be associated with HIV-1 control in lymphoid tissues. On the other hand, CD57 expression on CD8⁺ T cells was associated with increased tissue viral burden during early stages of ART and immunotherapy. Previous studies have shown that early KLRG1 expression, in the absence of CD57, is associated with CMV infection control ^74^ and lower proportion of CD57 on CD8+ T cells has been associated to enhanced antiviral responses in HIV-1 controllers ^33^. Therefore, it is plausible that the antibody could act at this level; however, further studies using this antibody in the context of HIV-1 infection are needed. Additionally, selective effects of TIGIT or KLRG1 blockade on NK cells or CD8+ T lymphocytes should also be addressed in future experiments through targeted depletion of each cell subset. Moreover, the potential latency-reversing or modulatory effect of aTIGIT- or aKLRG1-blocking mAbs on CD4+ T cells was not directly analyzed in our study and deserves further investigation. Previous studies have shown that latently HIV-1 infected cells express TIGIT ^75,76^ and may be reactivated upon TIGIT blockade, although in our previous study we observed variable reactivation in PWH ^28^ and additional studies are required to stablish the involvement of this receptor in viral latency. Similarly, *in vitro* studies have suggested that anti-KLRG1 mAbs can induce reactivation of HIV-1-infected CD4+ T cells ^31^. Therefore, in infected hBLT mice, these antibodies could exert a direct effect on HIV+ CD4+ T cells by reactivating latent virus, acting as latency-reversing agents (LRAs), and thereby promoting the elimination of infected cells either through cytopathic effects ^77^ or by exposing the virus to immune clearance. Additionally, the aTIGIT and aKLRG1 mAbs may influence homing of immune cells by modulating the expression of homing molecules such as CD44 which is expressed during T cell maturation as well as TIGIT ^78^, immune subset which could be targeted after treatment with aTIGIT mAb. These mechanisms should be assessed in settings where CD4+ T cells and other immune cells such as CD8+ T lymphocytes and NK cells are studied in isolation. Nevertheless, our current findings provide evidence suggesting differential regulation by these antibodies in the context of HIV infection in this humanized mouse model.

Another limitation of this model is that B cells do not mature properly and fail to generate secondary responses after repeated immunizations. As a result, fully developed secondary lymphoid organs, as seen in humans, do not form in these animals ^79^. Nevertheless, we were able to identify structures within secondary lymphoid tissues, such as the spleen, that resemble human white pulp. Within these white pulp-like areas, we observed a restricted localization of p24+ cells, which—as previously mentioned— could express KLRG1 ^31^, suggesting that they could be modulated by the binding of aKLRG1 mAb. In addition, IFNγ+ CD8+ T cells were also found within these structures near to infected cells specifically observed in the group of mice treated with aKLRG1 mAb, which could restrict the infected p24+ cells in this white pulp areas. Previous studies have described that CD8+ T cells tend to control infection around these viral niches but are typically unable to penetrate them ^80^. Therefore, we can speculate that this antibody may facilitate the entry or the migratory facility of IFNγ-producing CD8+ T cells into the lymphoid white pulp areas from these mice. In line with this, previous studies have identified follicular T cells expressing KLRG1+ with low levels of Bcl6, a transcription factor typically found in cell populations residing within lymphoid follicles ^81^, which may be associated with the maturation of these CD8+ T cells and their expression of CXCR5, facilitating their migration to B cell follicles. In contrast, in the hBLT mice treated with aTIGIT mAb, reduction of p24+ infected cells was not restricted by areas and observed in the whole tissue. The fact that other studies have shown that NKG2A+ NK cells selectively infiltrate lymph nodes ^82,83^, may be consistent with the observation that NKG2C+ cells do not enter the white pulp areas in mice treated with aTIGIT or aKLRG1 mAbs. Thus, viral control may be exerted outside of these areas by modulating the cytotoxic activity of these NKG2C+ cells, which express low level of NKG2A.

Surprisingly, a major finding of our study is the unexpected detrimental effect of the bispecific antibody, resulting in impaired viral control both in combination with ART and accelerated plasma viral load rebound following ATI. Our initial rationale was based on the hypothesis that simultaneous targeting of both TIGIT and KLRG1 exhaustion receptors would synergistically enhance immune function by reversing dysfunction in both NK cells and CD8+ T lymphocytes. This premise was based on previous observations of increased cytotoxic profiles of NK cells in response to K562 cell culture and in CD8+ T cell proliferation and activation induced by allogeneic DCs. However, *in vitro* evaluation after our *in vivo* analysis has revealed a specific increase in apoptosis marker Annexin V on NK cells upon treatment with the bispecific aTIGIT/aKLRG1 mAb, likely due to cellular overactivation. Alternatively, we could speculate that a fratricide effect mediated by ADCC from NK cells themselves may be occurring, as has been observed in treatments with Daratumumab ^84^. Another possibility is the fratricide effect of CD8+T cells in close proximity to NK cells, enhanced by bispecific mAb. These possibilities could explain the impaired viral control observed *in vivo*. Supporting the possibility of increased apoptosis in NK cells from hBLT treated with the bispecific mAb, we have observed lower proportions of CD16+ CD56dim and adaptive NKG2C+ NK cell in the blood and spleen of these mice. Surprisingly, the apoptosis was not observed in CD8+ T cell after treatment with the bispecific mAb *in vitro*. However, reduced proportions of activated and polyfunctional CD8+ T cells were observed *in vitro* in the presence of the bispecific mAb, suggesting that the function of these lymphocytes may be impaired. Therefore, besides confirming additional phenotypical and functional markers in tissue from hBLT treated with bispecific mAb, additional alternative dosing strategies or administration schedules should be explored to evaluate the therapeutic potential of this bispecific mAb and avoid the possible NK cell overactivation.

Finally, at the molecular level no differences were observed in the levels of integrated HIV-1 DNA in infected cells from lymphoid tissues once viral rebound had already occurred, with plasma viral loads approaching 10⁶ copies/mL across all groups. This made it difficult to detect differences at this stage. However, differences were clearly detected in infected lymphocytes CD4+ T cells from PB in aTIGIT and aKLRG1 mAb treated hBLT groups at the same time. Therefore, longitudinal assessment of proviral DNA levels in tissue should be conducted in this immunotherapy model to better understand the impact over time ^85^. Thus, despite these limitations, our study demonstrates that HIV-1 infection can be controlled in combination with ART and individual antibodies targeting either TIGIT or KLRG1, both at early time points and after ART interruption. Therefore, these immunotherapies could be further developed in future studies as potential strategies toward an HIV cure.

## DATA AVAILABILITY

This study includes no data deposited in external repositories.

## ACKNOWLEDGMENTS

The project leading to these results has received funding from “la Caixa” Banking Foundation under the project ETI-CURE code [LCF/PR/ HR20-00218]. E.M.G was supported by the Spanish Agencia Estatal de Investigación RETOS, Generación de conocimiento and consolidation programs (PID2021-127899OB-I00; CNS2023-144841), La Caixa Banking Fundation ETI-CureHIV (HR20-00218), GLD24/00117 grant from Gilead biosciences and infectious diseases CIBER from ISCIII (CB21/13/00107). M.A.L was supported by the Formación de Personal Investigador (FPI) grant PRE2022-104516. I.T. was supported by FPI UAM fellowship. R.G. was supported by by the ETI-Cure HR20-00218; PID2021-127899OB-I00 and GLD24/00117grants. P2022/BMD7209-INTEGRAMUNE from Comunidad Autónoma de Madrid and La Caixa Health Research Grant LCF/PR/HR23/52430018 and PID2023-149541OB-I00 to F.S.M also supported the study. I.S.C was supported by infectious diseases CIBER from ISCIII (CB21/13/00107). J.G.P was supported by La Caixa Health program ETI-CureHIV project (HR20-00218), the CIBERINFECC from the National Health Institute Carlos III, and the project PI22/01120. M.L.T was supported by the Spanish Agencia Estatal de Investigación, Ministerio de Ciencia, Innovación y Universidades (PID2022-138880OB-I00 and PDC2021-121238-I00). MJ-B is supported by the Miguel Servet program funded by the Spanish Health Institute Carlos III (CPII22/00005), and by the Spanish Agencia Estatal de Investigación, Ministerio de Ciencia, Innovación y Universidades (PID2021-123321OB-I00, PDC2022-133836-I00, and PID2024-155881OB-100).

## DICLOSURE AND COMPETING INTERESTS STATEMENT

The authors declare that they have no competing interests.

## SUPPLEMENTAL FIGURE LEGENDS

**Supplemental Figure 1.** I*n vitro* analysis of specificity and impact on functional NK cells of individual and bispecific antibodies directed to TIGIT and/or KLRG1. (A): Representative flow cytometry plots showing gating strategy of CD3+ T and CD3-CD56+ NK cells populations. Gated CD56dim CD16+ cells were selected within total CD56+ cells (B, C): Representative flow cytometry plots showing targeting of aTIGIT, aKLRG1 (B) or aTIGIT/aKLRG1 bispecific (C) mAbs on CD3+ T and CD3-CD56+ CD16+ NK cells populations through labeled secondary IgG1 antibody. An background with only secondary anti-IgG1 control mAb was included. (D-E): Dose response analysis of intracellular expression of CD107a and IFNγ in NK from a selected PWH after culture with increasing concentrations of aTIGIT (orange), aKLRG1 (green) and bispecific aTIGIT/aKLRG1 (purple) mAbs in the presence of K562 cells. (F): Fold change in proportions of CD107a+ IFNγ+ cells within CD56dim CD16+ NK cells from n=7 PWH, in response to the selected 5μg/ml of the different antibody treatment groups. (G): Fold change in proportions of CD107a+ IFNγ- and CD107a+ IFNγ+ cells within CD56+ CD16+ NK cells from n=7 PWH, in response to the selected 5μg/ml of bispecific aTIGIT/aKLRG1 mAb treated group. In panel D and E, data are presented as Box and Whiskers plots showing median values, with minimum and maximum error bars. Statistical significance was calculated using a using a Wilcoxon t test. *p<0.05.

**Supplemental Figure 2.** I*n vitro* functional analysis of individual and bispecific antibodies directed to TIGIT and/or KLRG1 in CD8+ T cells. (A, B): Representative flow cytometry plots showing gating strategy of CD8+ CD3+ T cells and CD107a versus IFNγ degranulation markers. (B): Representative flow cytometry plots showing CD107a versus TNFα in CD8+ T cells after targeting aTIGIT, aKLRG1 or aTIGIT/aKLRG1 bispecific mAbs in a MLR assay. An Isotype control mAb was included. (C): Fold change in proportions of CD107a+ TNFα+ cells within CD8+ T cells from n=6 PWH, in response to the selected 5μg/ml of the different antibody treatment groups. (D): Fold change in proportions of CD107a+ IFNγ- (left), CD107a+ IFNγ+ (center) and CD107a+ TNFα+ (right) cells within CD8+ T cells from n=6 PWH, in response to the selected 5μg/ml of bispecific mAb treated group. In panel C and D, data are presented as Box and Whiskers plots showing median values, with minimum and maximum error bars. Statistical significance was calculated using a using a Wilcoxon t test. *p<0.05.

**Supplemental Figure 3.** Improvement of NK cell reconstitution of hBLT mice by exogenous rhIL-15 administration and *in vivo* experimental design of HIV-1 infection, ART and immunotherapy. (A): Schematic representation of n=20 hBLT mice receiving three rhIL-15 injections every week. (B): Representative flow cytometry plots showing hCD45, hCD8, hCD56 and CD16 populations identifying NK cells after reconstitution of hBLT mice. (C): Absolute numbers of total hCD45+ as well as hCD4+, hCD8+ and hCD56+ cells in hBLT mice afer 1 and 2 weeks of rh-IL15 treatment. (D): Representative graphic of weights from different treated hBLT mice along the experiment. (E): Representative scheme of *in vivo* experiment showing HIV-1 infection, ART initiation and combination with either Isotype or aTIGIT or aKLRG1 or aTIGIT/aKLRG1 bispecific immunotherapy as well as ATI. Times of blood extraction, pVL quantification, proviral DNA analysis and histological studies are specified. (E): Weight of each individual hBLT mice during the duration of the *in vivo* experiment. Individual animals from the isotype, aTIGIT, aKLRG1 and bispecific mAb groups are labelled in blue, orange, green and purple, respectively. In panel C and E data are presented as Before-after plot showing individual values. Statistical significance was calculated using a using a Wilcoxon test. ****p<0.0001.

**Supplemental Figure 4.** Analysis of HIV-1 pVL levels before ART and suppression and viremia rebound after combination of ART with either individual or combined TIGIT or aKLRG1 blockade. (A): HIV plasma viral loads (pVL) prior to ART (8 days post-infection) comparing the different hBLT treated with isotype (blue), aTIGIT (orange), aKLRG1 (green) an aTIGIT/aKLRG1 bispecific (purple) mAb. (B): A Cox regression analysis showing time to achieve aviremia in each treatment antibody group during ART+IT therapy. (C): pVL of mice treated with bispecific aTIGIT/aKLRG1 mAb prior to ART initiation (D8) and during the two weeks of ART treatment (15 and 19 days post-infection) (left panel). Pie chart showing the percentage of viremic hBLT mice in bispecific mAb treatment group at day 15 (right panel). (D): Raw data plasma HIV-1 RNA levels at 6, 9, 12 and 16 days post-ATI compared to pre-ATI levels in isotype (blue), aTIGIT (orange), aKLRG1 (green) and bispecific (purple) groups. (E): Levels of pro-viral HIV-1 DNA normalized per 10^6^ CD4+ T cells in spleen from isotype (blue), aTIGIT (orange) and aKLRG1 (green) mAbs treated-hBLT-mice groups. (F): Fold change in plasma HIV-1 RNA levels at 6, 9, 12 and 16 days post-ATI compared to pre-ATI levels in bispecific (purple) group. (G): Levels of pro-viral HIV-1 DNA normalized per 10^6^ CD4+ T cells in PB from bispecific (purple) mAb treated-hBLT-mice group (left) and pie-chart showing the percentage of hBLT mice displaying integrated HIV-1 DNA values above or below the median values of the isotype hBLT group in bispecific mAb treated animal group (right). (H): Levels of pro-viral HIV-1 DNA normalized per 10^6^ CD4+ T cells in spleen from bispecific (purple) mAb treated-hBLT-mice group. (I): Representative confocal microscopy image showing expression of p24 (white) distributed in the spleen from an infected hBLT mouse. (J, K, L): Number of total p24+ cells (J), p24+ cells into WP (K), p24+ cells outside WP (L) normalized to total tissue area from isotype, aTIGIT and aKLRG1 treated mice groups. (M): Ratio between number of p24+ cells into white pulp areas and p24+ cells in total tissue from bispecific mice group. Pie charts showing proportions of animals below the median values of the isotype mAb control animal group are shown below. In panel A, C, D, E, F, G, H, J, K,L and M data are presented as Box and Whiskers or Violin plots showing median values, with minimum and maximum error bars. The red discontinued line respresents the median value of isotype-treated group. Pie chart data in panels C, G and K represent group proportions. Statistical significance was calculated using a using a Wilcoxon, Mann-Whitney test, and Chi-square test. *p<0.05; **p<0.01; ***p<0.001.

**Supplemental Figure 5.** Functional characterization of adaptive NKG2C+ NK cells in the spleen of HIV-1 infected hBLT mice treated with the different mAbs. (A): Representative flow cytometry plot showing gating of NKG2C and CD57 NK cells on human CD56dim CD16+ cells from hBLT mice. (B, C): Raw data reflecting proportions of NKG2C+ CD57- (C) and NKG2C-CD57+ (D) NK cells from isotype (blue), aTIGIT (orange) and aKLRG1 (green) mAb-treated hBLT mice groups (D): Ratio between NKG2C+ CD57- and NKG2C-CD57+ NK cells from isotype, aTIGIT and aKLRG1 BLT mice groups. (E, F): Raw proportions of CD56dim CD16+ (E), NKG2C+ CD57- (F, left) and NKG2C-CD57+ (F, right) NK cells from isotype (blue) versus bispecific aTIGIT/aKLRG1 mAb (purple) treated hBLT mice groups. (G): Ratio between NKG2C+ CD57- and NKG2C-CD57+ CD56dim CD16+ NK cells. (H): Representative flow cytometry plots of CD107a, IFNγ and TNFα in CD56dim CD16+ NK cells. (I, J): Spearman correlations between proportions of total CD107a+ cells (I) and CD107a+ IFNγ+ cells (J) within NKG2C+ CD57-NK cells versus HIV-1 pVL 16 days post-ATI in aKLRG1 (green) and isotype (blue) mAb-treated groups. (K): Spearman correlation between proportions of CD107a+ IFNγ+ cells in NKG2C-NK cells versus proportions of CD107a+ TNFα+ CD56+ CD16+ NK cells in the in aKLRG1 (green) and isotype (blue) mAb treated-hBLT mice. (L): Representative flow cytometry plots of NKG2C in CD3-CD56dim CD16+ NK cells and CD3+ CD8+. (M): Representative confocal microscopy image showing expression of NKG2C (green), p24 (white) and granzyme B (red) in the spleen from a representative isotype (left) and aTIGIT (right) mAbs treated hBLT mice. In panel B, C, D, E, F, G data are presented as Box and Whiskers showing median values, with minimum and maximum error bars. Statistical significance was calculated using a Mann-Whitney test or a Spearman correlation test. *p<0.05.

**Supplemental Figure 6.** Differential functional patterns of CD8+ T cells in HIV-BLT mice treated with individual aKLRG1 or aTIGIT mAbs and bispecific mAb. (A): Proportions of CD107a-IFNγ+ TNFα+ within CD8+ T cells in spleen from hBLT mice treated with an IgG isotype versus individual aTIGIT and aKLRG1 or bispecific aTIGIT/aKLRG1 mAbs. Pie-charts showing the percentage of hBLT mice displaying proportions of CD107a-IFNγ+ TNFα+ CD8+ T cells above or below the median values of the isotype hBLT group in the aTIGIT and aKLRG1 mAb treated animal groups are shown below plot in panel A. (C): Spearman correlation between proportions of CD107a-IFNγ+ TNFα+ CD8+ T cells in SP and fold change in HIV-1 pVL post-ATI in aTIGIT mAb treated hBLT mice. (D, E): Spearman correlation between proportions of CD107a- IFNγ+ TNFα+ CD8+ T cells and proportions of CD107a+ IFNγ+ NK cells (D) and proportions of CD107a+ IFNγ+ in NKG2C+ or NKG2C- NK cells (E) in SP from aTIGIT mAb hBLT treatment group. (F): Spearman correlation between proportions of CD107a+ CD8+ T cells in PB and proportions of CD107a+ TNFα+ CD56dim CD16+ NK cells in aKLRG1 group. Animals from aTIGIT, aKLRG1 and isotype mAb groups have been identified in orange, green and blue, respectively in panels C,D,E,F. (G): Spearman correlation between proportions of GZB+ CD8+ T cells in SP after ATI and HIV-1 pVL during ART combined with aKLRG1 mAb. (H): immunofluorescence histological analysis of IFNγ (green), HIV-1 p24 (white) and CD8 (red) in the spleen from a representative hBLT mouse treated with aTIGIT mAb. (I, J, K): Number of total CD8+ T cells (I), number of CD8+ IFNγ+ within WP (J) and outside WP (K), normalized to total tissue area in SP from the different hBLT treatment mice. In panel A, B, I, J and K data are presented as Box and Whiskers plots showing median values, with minimum and maximum error bars. Statistical significance was calculated using a Mann-Whitney, Chi-square and Spearman correlation test. *p<0.05; **p<0.01.

**Supplemental Figure 7.** Characterization of adaptive NKG2C+ NK cells in PB of HIV-1 infected hBLT mice treated with bispecific mAb at ART and immunotherapy initiation. (A, B, C): Proportions of CD56dim CD16+ NK cells (A) and proportions of NKG2C+ (B) and NKG2C- (C) cells within human CD56dim CD16+ NK cells in hBLT mice treated with IgG1 Isotypic (blue) and aTIGIT/aKLRG1 bispecific (purple) mAbs after ART initiation in combination with immunotherapy (IT) at 15 days post-infection. (D): Spearman correlation between proportions of CD57+ cells within circulating CD8+ T cells versus number of p24+ cells in the spleen normalized to total tissue area in aTIGIT (orange) and isotype (blue) mAb-treated hBLT group.

**Supplemental Figure 8.** Analysis of apoptosis induction in NK cells and CD8+ T cells by the aTIGIT/aKLRG1 mAb *in vitro*. (A, B): Proportions of Annexin V+ cells within NKG2C+ (A, left) and NKG2C- (A, right) CD56dim CD16+ NK cells and in CD8+ T cells (B).

