## Supplemental figures and tables for "Differential CD8+ T/NK cell-mediated reduction of HIV-1 replication after combination of ART with TIGIT or KLRG1 blockade in humanized mice"

### Supplemental Figure 1

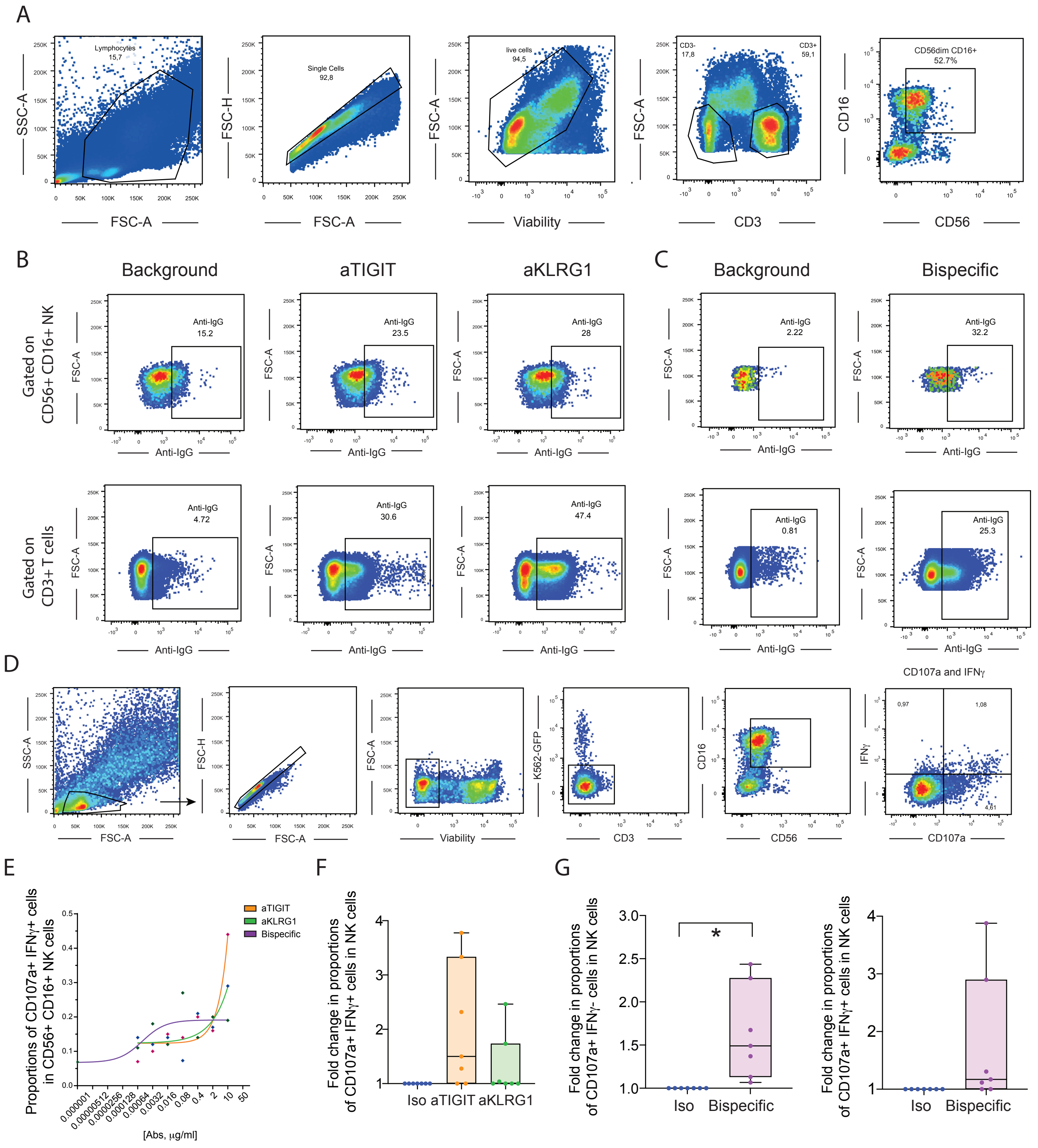

### Supplemental Figure 2

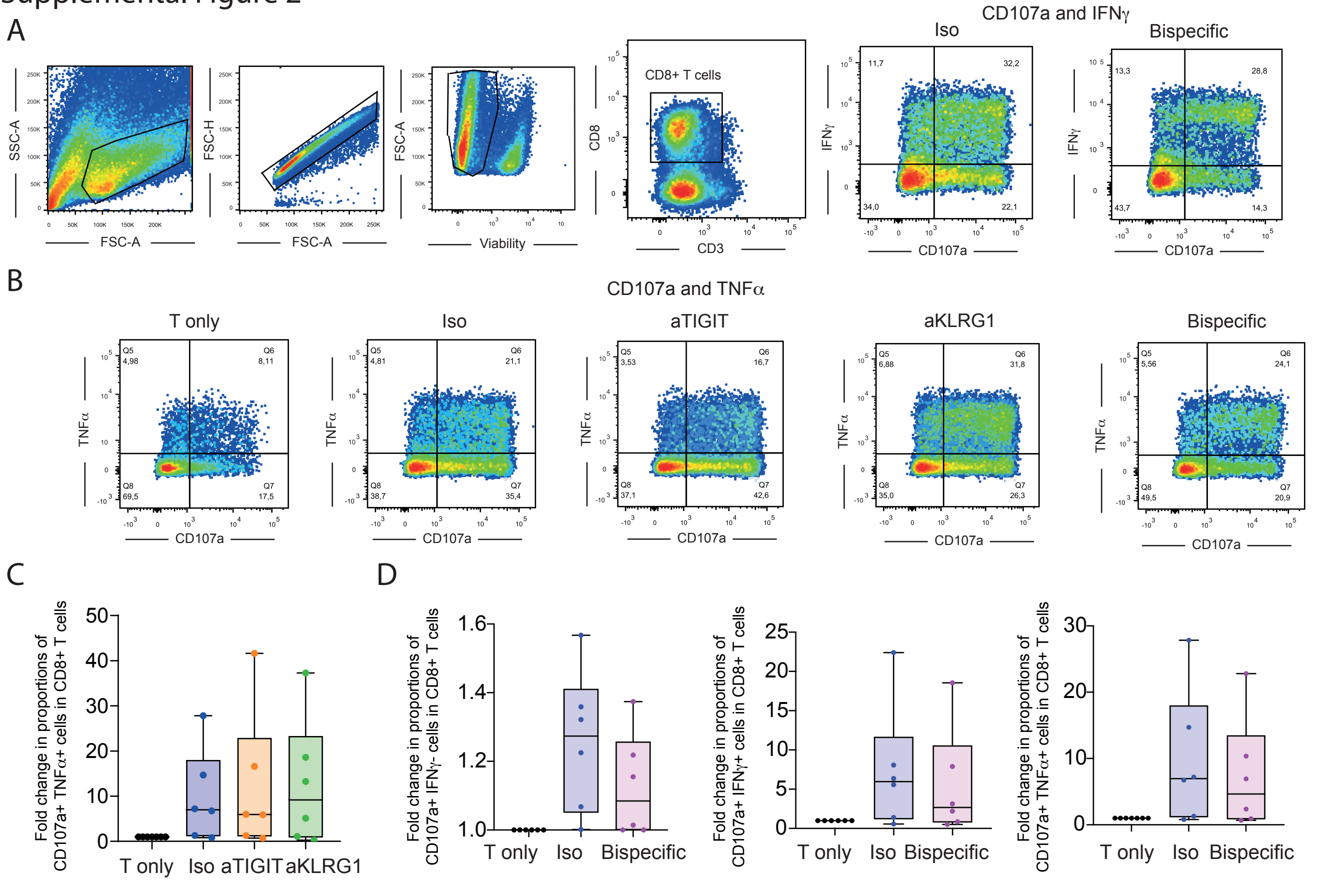

### Supplemental Figure 3

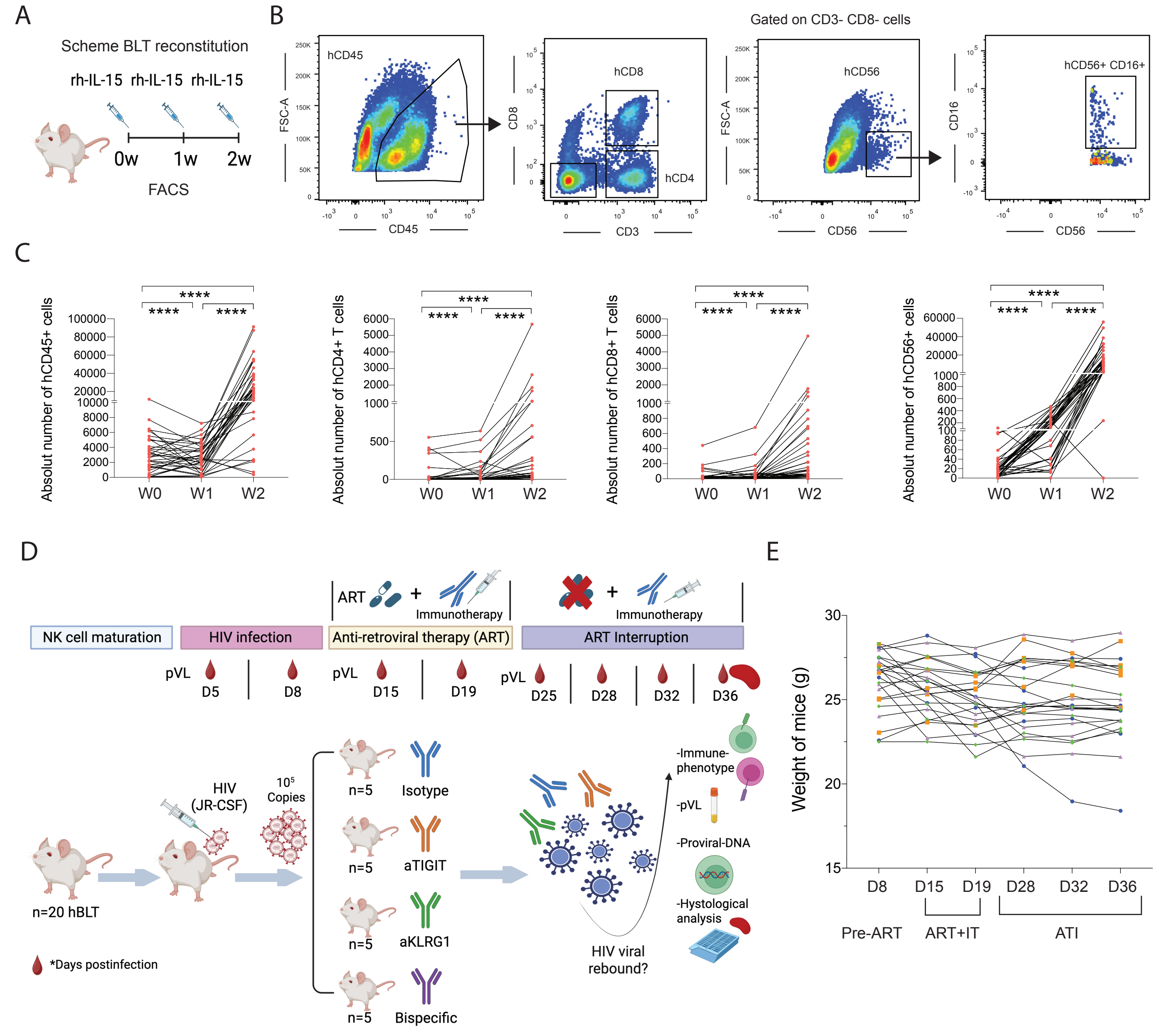

Supplemental Figure 4

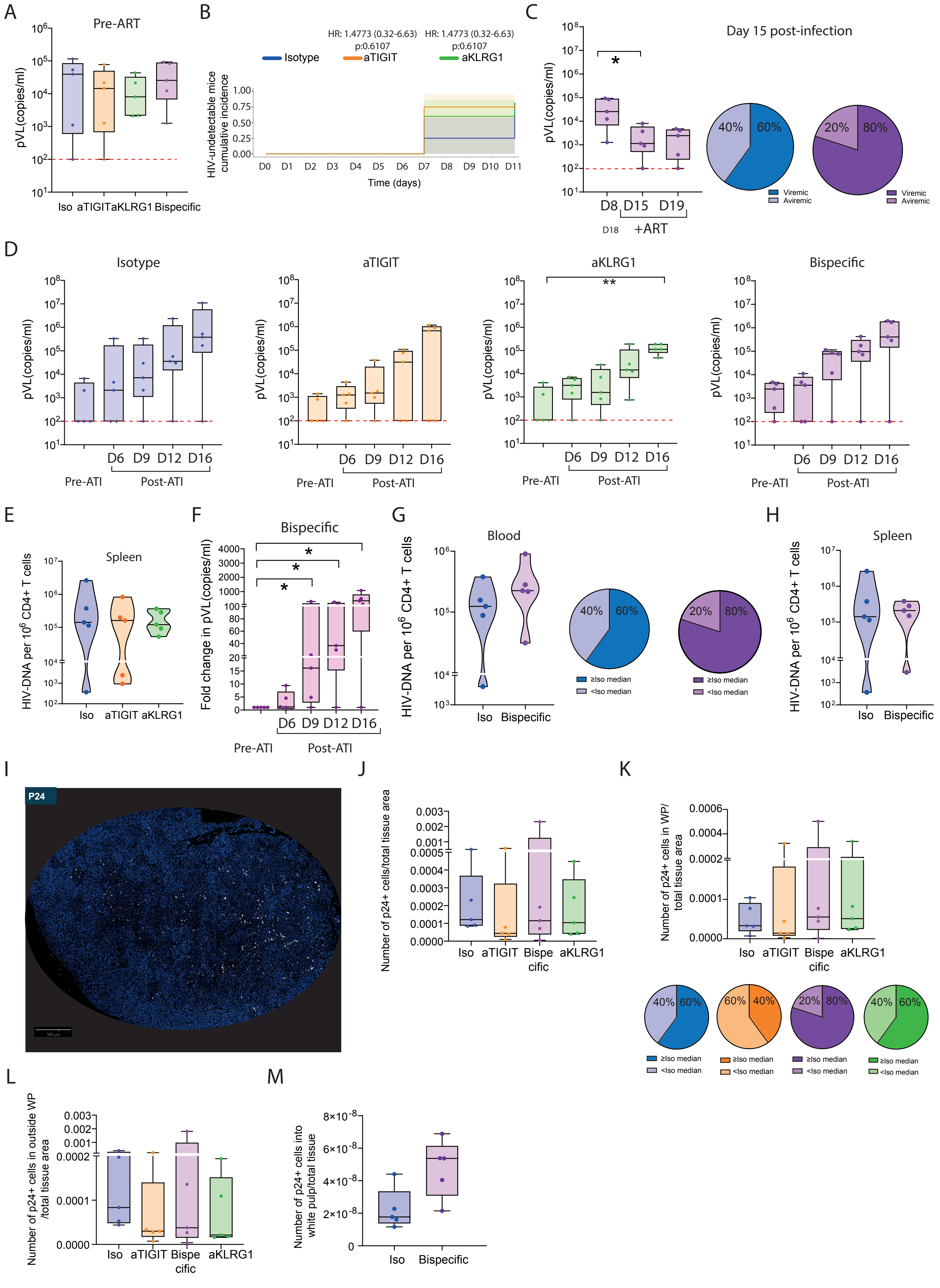

### Supplemental Figure 5

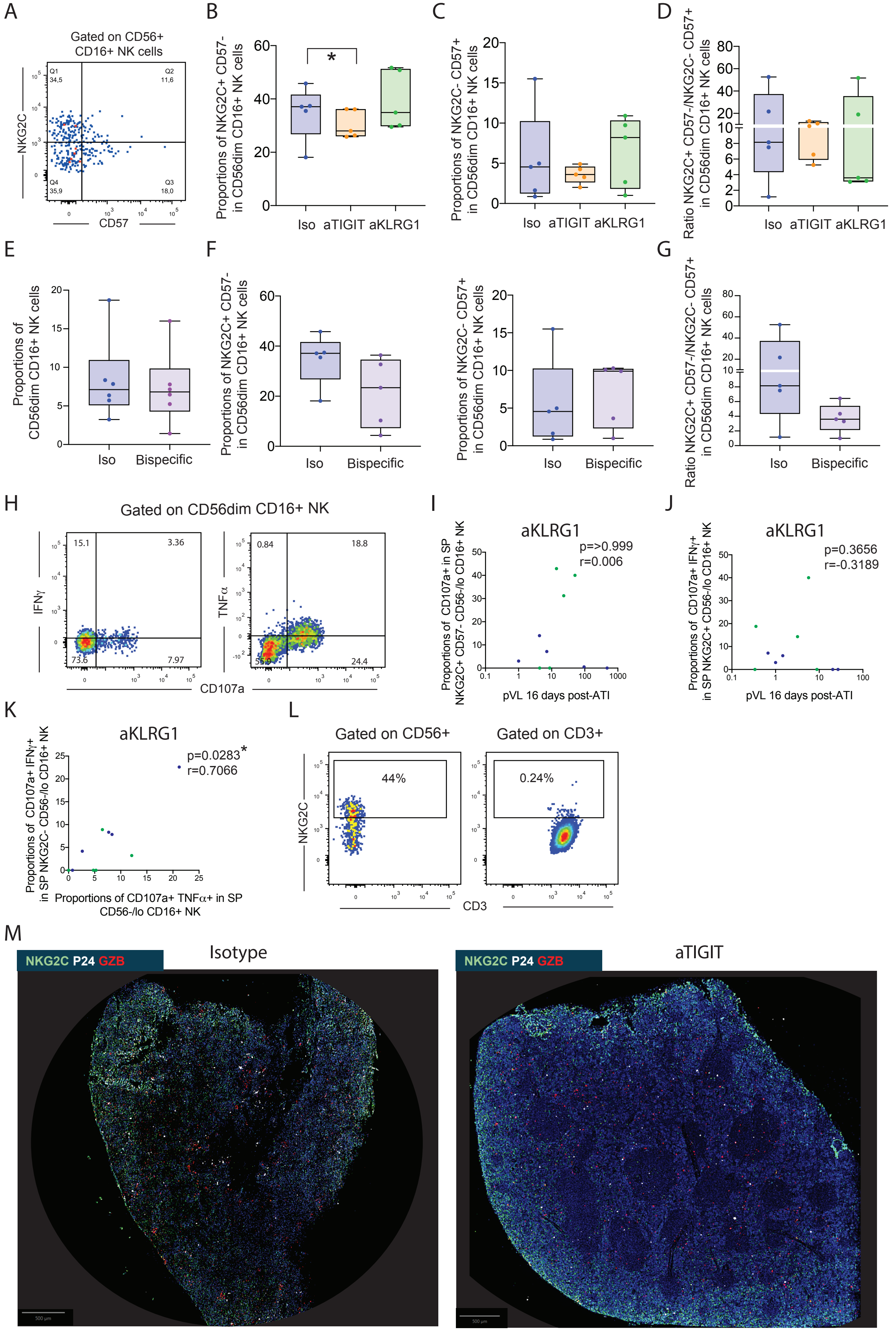

Supplemental Figure 6

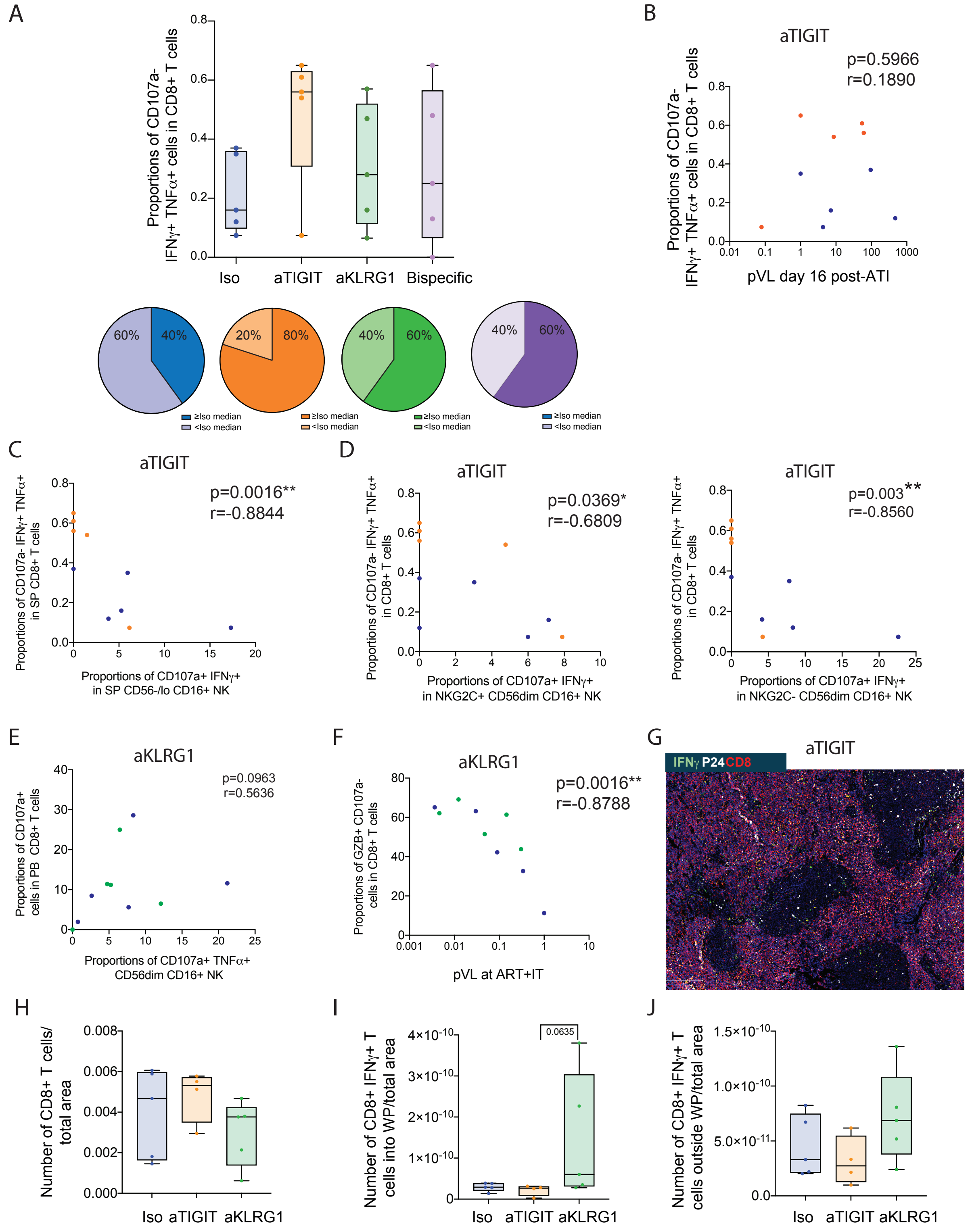

### Supplemental Figure 7

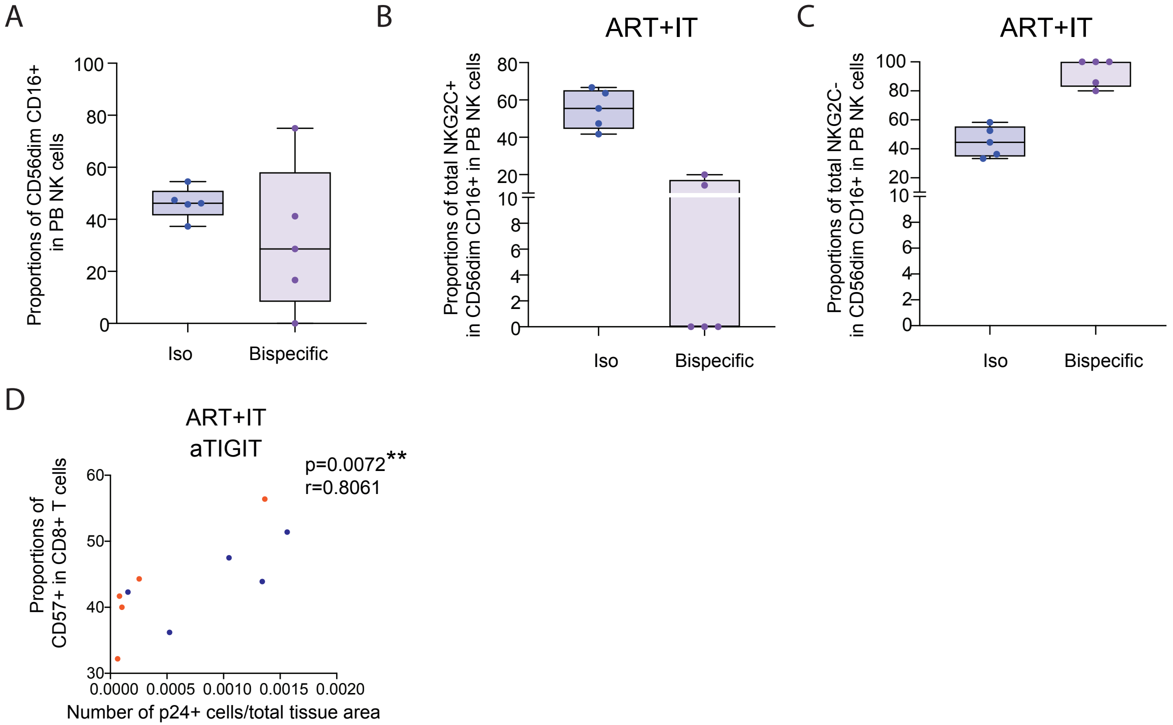

### Supplemental Figure 8

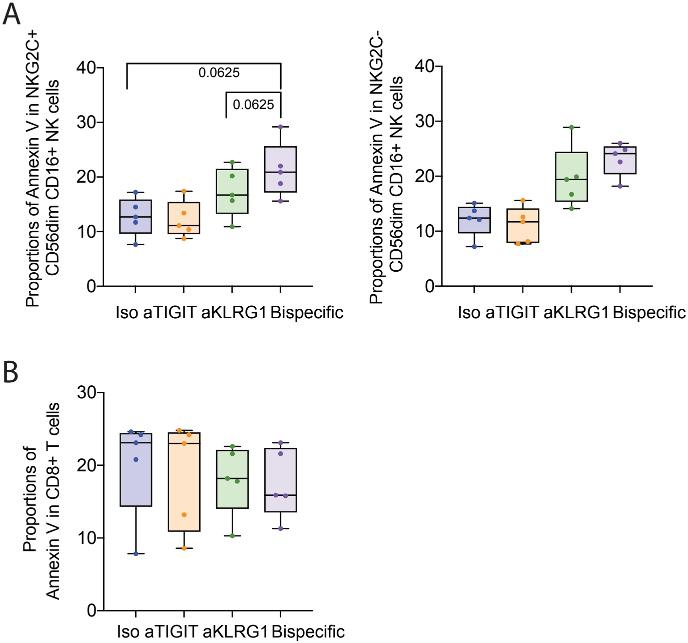

#### Supplemental Table 1. Reagents and Tools Table

| Reagent/Resource | Reference or Source | Identifier or Catalog Number |
|---|---|---|
| **Antibodies** | | |
| Anti-human CD3 | Becton-Dickinson (BD) | 565983 |
| Anti-human CD56 | Biolegend | 362505 |
| Anti-human CD8 | Becton-Dickinson (BD) | 612890 |
| Anti-human GZB | Becton-Dickinson (BD) | 563389 |
| Anti-human CD16 | Biolegend | 302045 |
| Anti-human CD45 | Becton-Dickinson (BD) | 560777 |
| Anti-human IgG1 | Biolegend | 410719 |
| Anti-human IFN$\gamma$ | Becton-Dickinson (BD) | 340452 |
| Anti-human TNF$\alpha$ | Biolegend | 502923 |
| Anti-human CD107a | Biolegend | 328620 |
| Anti-human CD56 | Biolegend | 362538 |
| Anti-human NKG2C | RyD Biotechne | FAB138G |
| Anti-human CD3 | Biolegend | 317334 |
| Anti-human CD16 | Biolegend | 302021 |
| Anti-human NKG2C | Biolegend | 375004 |
| Anti-human CD57 | Biolegend | 359623 |
| Anti-human CD8 | Biolegend | 300913 |
| Anti-human IFN$\gamma$ | Becton-Dickinson (BD) | 561980 |
| Ghost DyeTM | Tonbo Biosciences | 13-0865-T100 |
| Rat anti-human Granzyme B | Thermo Fisher | 14-8889-80 |
| Rabbit anti-human NKG2C | Abcam | AB230900 |
| Mouse anti-HIV-1 p24 | Abcam | Ab63958 |
| Rat anti-human CD8 | Bioi-Rad | MCA351GT |
| Rabbit anti-human IFN$\gamma$ | Abcam | Ab231036 |
| Donkey anti-rabbit AF488 | Thermo Fisher | R37118 |
| Goat anti-rabbit AF488 | Thermo Fisher | A-11008 |
| Donkey anti-rat AF594 | Jackson ImmunoResearch | 712-586-150 |
| Donkey anti-mouse AF647 | Thermo Fisher | A-31571 |
